# Rationally designed immunogens enable immune focusing to the SARS-CoV-2 receptor binding motif

**DOI:** 10.1101/2021.03.15.435440

**Authors:** Blake M. Hauser, Maya Sangesland, Kerri J. St. Denis, Ian W. Windsor, Jared Feldman, Evan C. Lam, Ty Kannegieter, Alejandro B. Balazs, Daniel Lingwood, Aaron G. Schmidt

**Affiliations:** Ragon Institute of MGH, MIT and Harvard, Cambridge, MA, 02139, USA; Department of Biological Chemistry and Molecular Pharmacology, Harvard Medical School, Boston, MA 02115, USA; Laboratory of Molecular Medicine, Boston Children’s Hospital, Boston, MA, USA; Department of Microbiology, Harvard Medical School, Boston, MA 02115, USA

**Keywords:** immunogen design, glycan, immune focusing, SARS-CoV-2, coronavirus

## Abstract

Eliciting antibodies to surface-exposed viral glycoproteins can lead to protective responses that ultimately control and prevent future infections. Targeting functionally conserved epitopes may help reduce the likelihood of viral escape and aid in preventing the spread of related viruses with pandemic potential. One such functionally conserved viral epitope is the site to which a receptor must bind to facilitate viral entry. Here, we leveraged rational immunogen design strategies to focus humoral responses to the receptor binding motif (RBM) on the SARS-CoV-2 spike. Using glycan engineering and epitope scaffolding, we find an improved targeting of the serum response to the RBM in context of SARS-CoV-2 spike imprinting. Furthermore, we observed a robust SARS-CoV-2-neutralizing serum response with increased potency against related sarbecoviruses, SARS-CoV, WIV1-CoV, RaTG13-CoV, and SHC014-CoV. Thus, RBM focusing is a promising strategy to elicit breadth across emerging sarbecoviruses and represents an adaptable design approach for targeting conserved epitopes on other viral glycoproteins.

**One Sentence Summary:** SARS-CoV-2 immune focusing with engineered immunogens

## MAIN TEXT

Humoral responses elicited by vaccination or infection predominantly target surface-exposed viral glycoproteins. These responses can often provide protection against future infections to the same or closely related viral variants. However, in some instances, such as influenza and HIV, the elicited responses are often poorly protective as they target variable epitopes (*1, 2*). Furthermore, waning of responses (*i*.*e*., durability), as is the case for common cold-causing coronaviruses, results in susceptibility to reinfections (*3-6*). For SARS-CoV-2 (SARS-2) it remains unclear whether current vaccines will confer long-term protection. Furthermore, it is increasingly apparent that humoral immunity elicited by vaccination or natural infection may provide reduced protection against emerging SARS-2 variants (*7-12*). Thus, implementing rational design strategies aimed at directing the immune response to conserved viral epitopes may help reduce the likelihood of viral escape and lead to more broadly protective responses (*13, 14*).

Two immunogen design strategies used to direct humoral responses include “masking” epitopes via engineering putative N-linked glycosylation sites (PNGs) and the design of protein scaffolds to present broadly protective epitopes (*15, 16*); these strategies have been used previously for viral glycoproteins RSV F, influenza hemagglutinin and HIV envelope (*17-19*). Applying these approaches to the SARS-2 spike provides an opportunity to potentially improve serum neutralization potency, efficacy against variants, and cross-reactivity of antibody responses. A potential target of these efforts is the angiotensin converting enzyme 2 (ACE2) receptor binding motif (RBM) of the receptor binding domain (RBD) (*20, 21*). Indeed, several potently neutralizing RBM-directed antibodies that interfere with ACE2 binding are protective and some can also neutralize related sarbecoviruses (*13, 14, 21-23*). Here, we show that hyperglycosylation of the RBD and a “resurfacing” approach that grafts the RBM from SARS-2 onto heterologous coronavirus RBDs focuses serum responses to the RBM. This immune-focused response is potently neutralizing with breadth across SARS-2 variants and other coronaviruses.

The RBM of SARS-2 and related sarbecoviruses, SARS-CoV (SARS-1) and WIV1-CoV (WIV1), is a contiguous sequence spanning residues 437-507 (SARS-2 numbering) of the spike protein. In an effort to elicit RBM-specific responses only, we first asked whether the RBM itself could be recombinantly expressed in absence of the rest of the RBD (**Fig. 1A**). While the SARS-2 RBM could indeed be overexpressed, it failed to both engage the conformationally-specific RBM-directed antibody B38 and bind to cell-surface expressed ACE2 (**Fig. S1**). These results likely suggest that the RBM is conformationally flexible, and that the RBD serves as a structural “scaffold” to stabilize the RBM in its binding-compatible conformation. To circumvent the considerable hurdle of *de novo* scaffold design for RBM presentation, we asked whether heterologous sarbecovirus RBDs from SARS-1 and WIV1 and the more distantly related merbecovirus MERS-CoV (MERS) could serve as scaffolds (**Fig. 1A**) —variations of this approach were used previously to modulate ACE2 binding properties (*24, 25*). In context of immunizations, we hypothesized that these heterologous RBDs would present the SARS-2 RBM while removing any other SARS-2-specific epitopes. The SARS-1, WIV1 and MERS RBDs share a pairwise amino acid identity with SARS-2 of 73.0%, 75.4% and 19.5%, respectively. The RBM is less conserved despite have a shared ACE2 receptor for SARS-1 and WIV1 with only 49.3% and 52.1% identity, respectively; as MERS uses DPP4 as a receptor, its RBM shares no notable identity (*26*). While we were unable to “resurface” MERS RBD with the SARS-2 RBM, the related SARS-1 and WIV1 RBDs successfully accepted the RBM transfer. These resurfaced constructs, rsSARS-1 and rsWIV1 retained binding to the SARS-2 RBM-specific B38 antibody as well as effectively engaged ACE2 (**Fig. S2**) (*22*). These data suggest that there are sequence and structural constraints within the RBD required for successful RBM grafting; such an approach may be facilitated by using CoV RBDs that use the same receptor for viral entry.

**Fig. 1.**
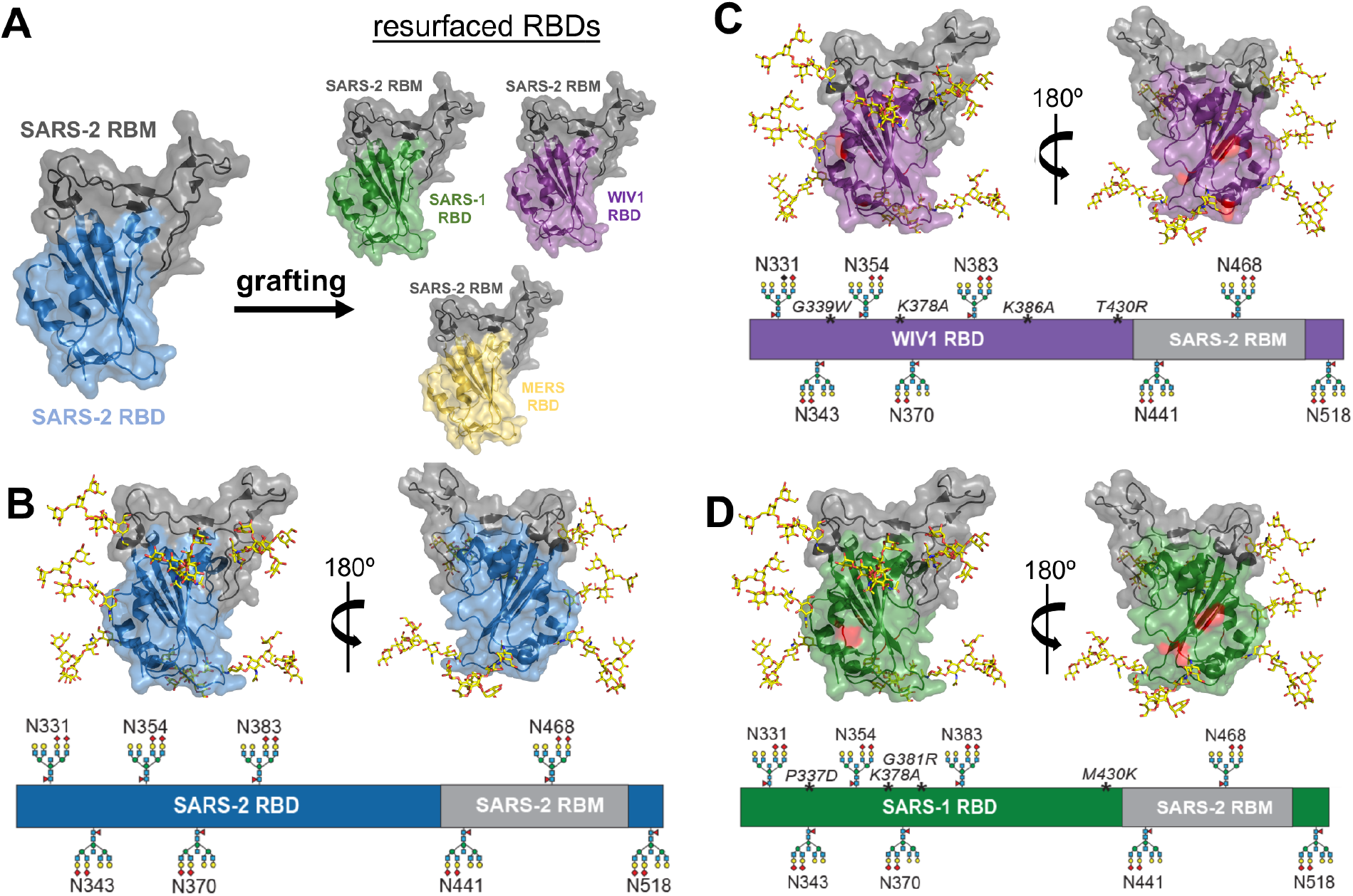
Resurfacing and hyperglycosylation approaches for immune-focusing. **(A)** Design schematic for resurfacing SARS-1 (rsSARS-1) and WIV1 (rsWIV1) with the SARS-2 receptor binding motif (RBM). Design schematic for hyperglycosylating SARS-2 **(B)**, rsSARS-1 **(C)** and rsWIV1 **(D)** receptor binding domains (RBDs). Non-native engineered glycans and native glycans are modeled; native SARS-2 RBM glycan at position 331 is omitted in the schematic. Mutations in the WIV1 and SARS-1 RBDs are shown in red and italicized in the linear diagram. All images were created using PDB 6M0J.

We next used these resurfaced RBDs as templates for further modification using glycan engineering. This approach aimed to mask conserved, cross-reactive epitopes shared between the SARS-1, SARS-2, and WIV1 RBDs. There are two evolutionarily conserved PNGs at positions 331 and 343; SARS-1 and WIV1 have an additional conserved PNG at position 370 (SARS-2 numbering). To further increase overall surface glycan density, we introduced novel PNGs onto wildtype SARS-2 as well as rsSARS-1 and rsWIV1 RBDs. Based on structural modeling and biochemical validation, we identified 5 potential sites on rsSARS-1 and rsWIV1 as well as 6 on SARS-2. Including the native PNGs, all constructs had a total of 8 glycans (**Fig. 1B-D, S3-4**)— we denote these hyperglycosylated (hg) constructs as SARS-2^*hg*^, rsSARS-1^*hg*^, and rsWIV1^*hg*^. We expressed these constructs in mammalian cells to ensure complex glycosylation in order to maximize any glycan “shielding” effect. We subsequently characterized these constructs using the

RBM-directed antibody B38, as well as ACE2 binding, to ensure that the engineered PNGs did not adversely affect the RBM conformation. Overall, the hyperglycosylated constructs were largely comparable in affinity for B38, with only ∼2-fold decrease, and still effectively engaged ACE2 (**Fig. S5**). These results confirm a conformational and functionally intact RBM

Next, we assessed whether the engineered PNGs abrogated binding to sarbecovirus cross-reactive antibodies S309 and CR3022—both antibodies were isolated from SARS-1 convalescent individuals (*27, 28*). The CR3022 contact residues on SARS-1 and WIV1 differ only at a single residue while SARS-2 differs at 5 residues across both CR3022 and S309 epitopes (*29*). Importantly, these epitopic regions were shown to be a significant portion of the SARS-2 RBD-directed response in murine immunizations and thus any RBM focusing would require masking of these regions (**Fig. S5**) (*27, 28, 30*). While SARS-2^*hg*^ effectively abrogated S309 and CR3022 binding, the engineered PNGs at the antibody:antigen interface on rsSARS-1^*hg*^ and rsWIV1^*hg*^ did not completely abrogate S309 and CR3022 binding. We therefore incorporated unique mutations on rsSARS-1^*hg*^ and rsWIV1^*hg*^ so that any elicited antibodies would be less likely to cross-react between these two constructs. To that end, we found K378A and the engineered glycan at residue 383 (SARS-2 numbering) completely abrogated CR3022 binding in both rsSARS-1^*hg*^ and rsWIV1^*hg*^ (**Fig. S5**). For S309, mutations P337D in rsSARS-1^*hg*^ and G339W in rsWIV1^*hg*^ in addition to glycans at residues 441 and 354 (SARS-2 numbering) were sufficient to disrupt binding (**Fig. S5**). We made two additional mutations, G381R, M430K on rsSARS-1^*hg*^ and K386A, T430R on rsWIV1^*hg*^, to further increase the antigenic distance between these scaffolds (**Fig. 1C, D**).

We then tested the immunogenicity and antigenicity of our optimized constructs and assessed their RBM immune-focusing properties, in the murine model. In order to increase avidity and to minimize any off-target tag-specific responses, we generated trimeric versions of each immunogen using our previously characterized hyperglycosylated, cysteine-stabilized GCN4 tag (*hg*GCN4^cys^) (*30, 31*). We first primed all cohorts with SARS-2 spike to reflect pre-existing SARS-2 immunity and to imprint an initial RBM response that may be recalled and selectively expanded by our immunogens. To test potential RBM immune-focusing, one cohort was sequentially immunized with SARS-2^*hg*^ trimers (“Trimer^*hg*^ cohort”) and a second cohort was immunized with SARS-2^*hg*^ trimers followed by a cocktail of rsSARS-1^*hg*^ and rsWIV1^*hg*^ (“Cocktail^*hg*^ cohort”) (**Fig. 2A**). In order to compare the efficacy of RBM-focusing, we included a “ΔRBM cohort”. This cohort was immunized with a modified SARS-2 RBD (ΔRBM) with four novel glycans engineered at positions 448, 475, 494, and 501 on the RBM. These PNGs effectively abrogate RBM-directed B38 antibody binding and engagement of ACE2 (*30*) and should restrict elicited humoral responses to this epitope. Finally, as a control cohort, we included a SARS-2 spike prime followed with sequential immunizations with wildtype (*i*.*e*., unmodified) SARS-2 RBD trimer (“Trimer cohort”).

**Fig. 2.**
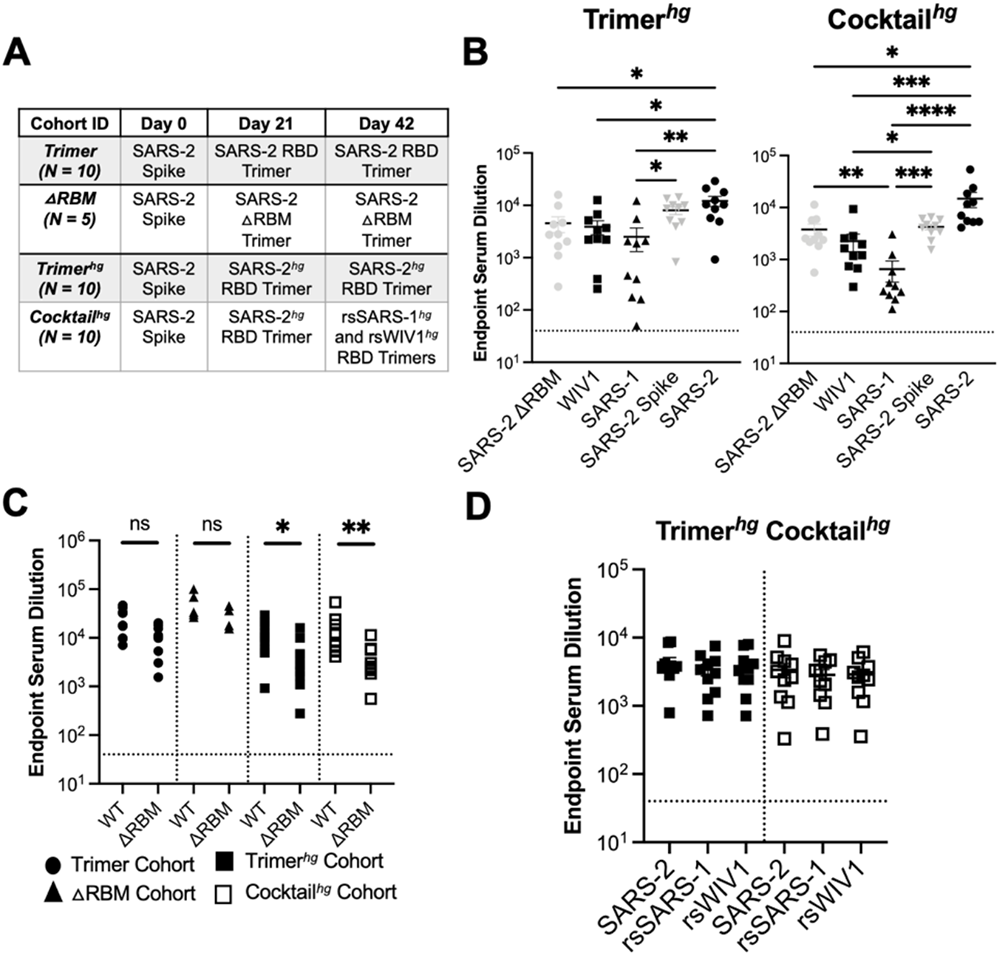
Serum analysis from cohorts. **(A)** Schematic of immunization cohorts; N= number of mice in each cohort **(B, C)** Serum following immunizations was assayed in ELISA at day 56 with different coronavirus antigens. Statistical significance was determined using Kruskal-Wallis test with post-hoc analysis using Dunn’s test corrected for multiple comparisons or Mann-Whitney U test (* = p < 0.05, ** = p < 0.01, *** = p < 0.001, **** = p < 0.0001). **(D)** Day 56 serum samples assayed against rsSARS-1 and rsWIV1 RBDs no longer show statistically significant differences in binding compared to SARS-2 RBD as determined using Kruskal-Wallis test with post-hoc analysis using Dunn’s test corrected for multiple comparisons.

Overall, we find that all cohorts elicit robust serum responses to wildtype SARS-2 RBD (**Fig. 2B-C, S5**). In order to specifically evaluate the RBM-directed responses, we compared serum ELISA titers to wildtype SARS-2 RBD and the SARS-2 ΔRBM RBD construct. We find that the Trimer^*hg*^ and Cocktail^*hg*^ cohorts had a significant increase in serum titers to wildtype SARS-2 RBD relative to SARS-2 ΔRBM RBD; this was in contrast to the ΔRBM and Trimer cohorts (**Fig. 2B,C, S6A**,**B**). Across the Trimer^*hg*^ and Cocktail^*hg*^ cohorts, the mean binding loss to the SARS-2 ΔRBM RBD relative to wildtype SARS-2 RBD was 64%, indicating that ∼64% of serum antibodies are RBM-directed by this metric (It is possible, however, that our engineered glycans ΔRBM construct does not fully restrict access by all RBM-directed responses with differing angles of approach). The Cocktail^*hg*^ cohort had a slight increase in RBM focusing relative to the Trimer^*hg*^ cohort. This may be due to increasing the overall antigenic distance (*i*.*e*., sequence difference) between the WIV1 and SARS-1 RBDs relative to SARS-2 while maintaining the identical SARS-2 RBM epitope. Additionally, we find that the Trimer^*hg*^ and Cocktail^*hg*^ cohorts had significantly lower titers to SARS-1 and WIV1 RBDs as compared to SARS-2 RBD (**Fig. 2B**). This difference was most pronounced in the Cocktail^*hg*^ cohort, suggesting that the hyperglycosylation and engineered mutations within the RBD effectively dampened responses to these conserved, cross-reactive epitopes that are present outside the RBM. Furthermore, serum titers against the rsSARS-1 and rsWIV1 RBDs were comparable to SARS-2 RBD, indicating that there is minimal antibody response directed towards wildtype SARS-1 and WIV1 RBD epitopes in comparison to the SARS-2 RBM (**Fig. 2D, S6C**). We observed no significant glycan-dependent serum response in either cohort that used hyperglycosylation (**Fig. S7**). Collectively, these data confirm an enhanced SARS-2 RBM-focused serum response elicited by our engineered immunogens.

We next compared the neutralization potency of all cohorts using SARS-1, SARS-2, and WIV1 pseudoviruses, as well as RaTG13-CoV (RaTG13) and SHC014-CoV (SHC014) (*25, 32-34*). While all cohorts elicited a potent SARS-2 neutralizing response, notably, the Trimer^*hg*^ and Cocktail^*hg*^ cohorts also exhibited potent SARS-1, WIV1, RaTG13, and SHC014 pseudovirus neutralization relative to the Trimer and ΔRBM cohorts (**Fig. 3A, S8-9, S6B**,**D**). This is particularly noteworthy for the Trimer^*hg*^ cohort as it did not include SARS-1 or WIV1 RBDs in the immunization regimen. WIV1, RaTG13, and SHC014 in this instance are broadly representative of possible future emerging sarbecoviruses with pandemic potential (*25, 34, 35*). The Trimer cohort lost significant neutralization against RaTG13, SARS-1, WIV1, and SHC014, and the ΔRBM trended towards a loss in neutralization as well, mirroring patterns seen following SARS-2 infection and immunization in humans, as well as SARS-2 spike-based immunization in mice (*8, 32, 36*). While the Trimer^*hg*^ cohort had a significant loss in neutralization against the most genetically divergent sarbecovirus, SHC014, it retained potency against RaTG13, SARS-1, and WIV1. Importantly, the Cocktail^*hg*^ had no significant loss in neutralization against either RaTG13 or SHC014, neither of which are vaccine-matched strains (**Fig. 3B, S8-9**).

**Fig. 3.**
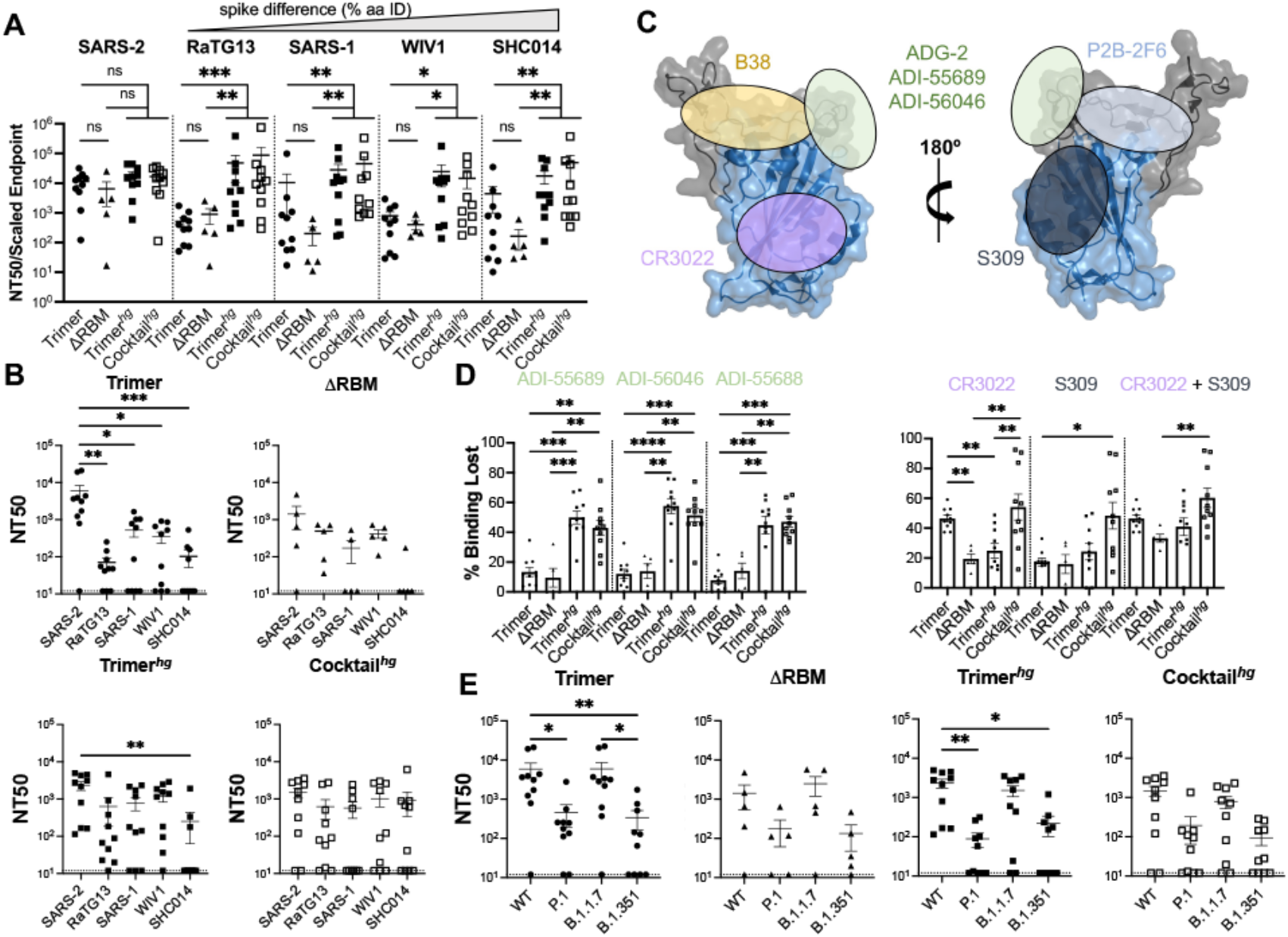
Potency and characterization of SARS-like coronavirus neutralization response. **(A)** Day 56 serum from all mice was assayed for neutralization against SARS-2, RaTG13, SARS-1, WIV1, and SHC014 pseudoviruses (arranged in order of genetic similarity of the full-length spike to SARS-2). Neutralization potency was computed using scaled endpoint serum ELISA titers. Statistical significance was determined using the Kruskal-Wallis test with post-hoc analysis using Dunn’s test corrected for multiple comparisons (* = p < 0.05, ** = p < 0.01, *** = p < 0.001, ns = not significant). **(B)** Day 56 serum from all mice was assayed for neutralization against SARS-2, RaTG13, SARS-1, WIV1, and SHC014 pseudoviruses. Statistical significance was determined using the Kruskal-Wallis test with post-hoc analysis using Dunn’s test corrected for multiple comparisons (* = p < 0.05, ** = p < 0.01, *** = p < 0.001). **(C)** Approximate locations of representative antibody epitopes from each of the four SARS-2 RBD-directed antibody classes (*21*) and ADG-2-like antibodies on the SARS-2 RBD. (PDB: 6M0J) **(D)** Antibody competition ELISAs with WIV1 RBD as the coating antigen. Bars show the mean percent binding lost, with error bars representing the standard error of the mean. Comparisons were performed using the Kruskal-Wallis test with post-hoc analysis using Dunn’s test corrected for multiple comparisons (* = p < 0.05, ** p < 0.01, *** = p < 0.001, **** = p < 0.0001). **(E)** Day 56 serum was assayed against SARS-2 variant pseudoviruses for neutralization. Statistical significance was determined using the Kruskal-Wallis test with post-hoc analysis using Dunn’s test corrected for multiple comparisons (* = p < 0.05, ** = p < 0.01).

To further epitope map the RBM-focused responses, we performed ELISA-based antibody competition using cross-reactive antibodies CR3022, S309, ADI-55688, ADI-55689, and ADI-56046 and WIV1 RBD (**Fig. 3C-D**). The latter two antibodies bind a conserved sarbecovirus RBM epitope also targeted by the antibody ADG-2, which is currently in clinical development and for which ADI-55688 is a precursor, and other antibodies with broad sarbecovirus neutralization (*13, 14, 37*). Competition ELISAs suggest that the cross reactive WIV1-directed responses in the Trimer^*hg*^ and Cocktail^*hg*^ cohorts focus to the ADG-2-like epitope, as well as to the CR3022 and S309 epitopes in the Cocktail^*hg*^ cohort (**Fig. 3C-D**). Thus, SARS-2^*hg*^, rsSARS-1^*hg*^, and rsWIV1^*hg*^ RBDs can induce not only potent SARS-2 neutralizing antibodies, but also cross-reactive antibodies that bind to a conserved RBM epitope (**Fig. S10**). Notably, these results are in contrast to our previous work showing that a cocktail of sarbecovirus that included SARS-1 and WIV1 RBDs could predominantly focus the antibody response towards the conserved CR3022 and S309 epitopic regions (*30*).

Many SARS-2 variants of concern include mutations within the RBM including B.1.1.7, B.1.351 and P.1 first detected in the United Kingdom, South Africa, and Brazil, respectively (**Fig. S11A**). We therefore asked what the consequence was of enhanced focusing to the RBM and whether the elicited responses elicited were sensitive to these mutations. Interestingly, serum from the Cocktail^*hg*^ cohort showed no significant loss of binding to the B.1.351 RBD compared to the wildtype SARS-2 RBD (**Fig. S11B**). This is in contrast to the control Trimer cohort and the Trimer^*hg*^ cohort, which showed a significant loss of binding and parallels the observation of reduced serum binding from human subjects immunized with current SARS-2 vaccines (*8, 38, 39*). Second, we tested all sera for neutralization against SARS-2 variant pseudoviruses: B.1.1.7, B.1.351 and P.1. While the control Trimer cohort and the Trimer^*hg*^ cohort could still neutralize all pseudoviruses, there was a significant loss of neutralization to the P.1 and B.1.351 variants, consistent with our ELISA data. In contrast, we find no significant loss of neutralization against these variants in the Cocktail^*hg*^ and ΔRBM cohorts (**Fig. 3E**). For ΔRBM cohort, the elicited responses were likely focused to neutralizing epitopes within the non-RBM RBD (*e*.*g*., CR3022, S309) and therefore were not sensitive to these RBM mutations. However, the neutralizing response observed in the Cocktail^*hg*^ cohort potentially indicates that substantial immune-focusing to the RBM may allow for greater recognition (i.e., accommodation) of mutations compared to the RBM-directed antibody response elicited via infection or vaccination (*38, 40*).

We next isolated SARS-2 RBD-specific IgG+ B cells from the Trimer, Trimer^*hg*^, and Cocktail^*hg*^ cohorts and obtained paired heavy and light chain sequences (**Fig. S12**). Overall, there was a predominance of IGHV1-42 gene usage across all cohorts, but light chain usage patterns varied more noticeably between the control Trimer cohort and the Trimer^*hg*^ and Cocktail^*hg*^ cohorts (**Fig. 4A**). CDRH3 length was significantly longer in the Trimer^*hg*^ cohort; mean somatic hypermutation trended towards being higher in the Trimer^*hg*^ and Cocktail^*hg*^ cohorts compared to the Trimer cohort (**Fig. 4B-C**). We recombinantly expressed representative antibodies from clonally related populations from the Trimer^*hg*^ and Cocktail^*hg*^ cohorts to test for breadth and to crudely epitope map (**Fig. 4D, S13; Table S1**). Ab19 and Ab20 were SARS-2 specific, did not bind the ΔRBM construct and did not compete with CR3022, suggesting an RBM-focused epitope. Importantly, Ab15, Ab16, and Ab17 were exceptionally broad in their reactivity, engaging all coronavirus RBDs tested as well as the SARS-2 variant B.1.351 (**Fig. 4D**). These antibodies still bound the ΔRBM construct and either completely (Ab15) or partially (Ab16, Ab17) competed with CR3022; affinities to the B.1.351 and ΔRBM construct were between ∼2-20 fold lower than the affinity to the SARS-2 RBD. These data suggest a conserved epitope that partially overlaps both the CR3022 epitope and the RBM (**Fig. S10**).

**Fig. 4.**
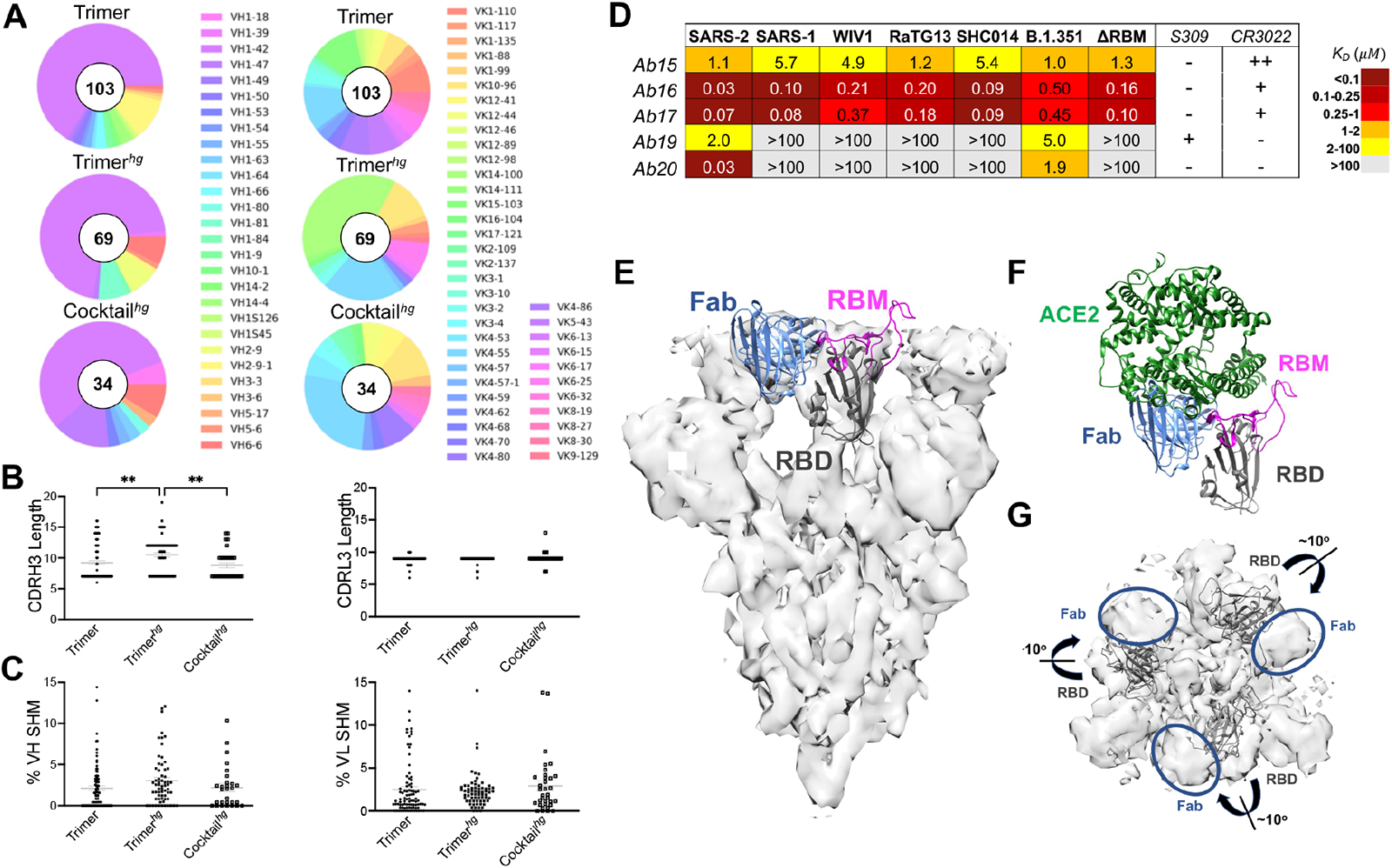
SARS-2 RBD-directed B cell characteristics. Spleens were harvested at day 63 and SARS-2 RBD-directed IgG+ B cells were isolated via flow cytometry. B cell receptor sequencing was used to characterize (**A**) heavy and light chain V-gene usage. All gene families listed are *01 except VH1-84*02. Complementarity determining region 3 (CDR3) length **(B)** and percent somatic hypermutation (SHM) **(C)** were also analyzed for each sequence. SHM was not analyzed for cohorts with uncertain IMGT V-gene assignments. Statistical significance was determined using the Kruskal-Wallis test with post-hoc analysis using Dunn’s test corrected for multiple comparisons (* = p < 0.05, ** = p < 0.01). **(D)** Antibodies representative of lineages that were expanded in RBM-focusing cohorts were expressed recombinantly as Fabs, and their binding was characterized via BLI (**E**) low resolution cryoEM map with model of Ab16 as Fab (blue) bound to RBD (gray) with the RBM (magenta) shown (RBD is from PDB 7DX9); for ease of viewing only a single RBD and Fab are shown. (**F**) Model from (**E**) with docked ACE-2 (from PDB 6M0J) (**G**) cryoEM map with 3 RBDs (gray) in ribbon and the Fab Ab16 removed to show its density and the slight outward rotation of the RBD required to better fit the density compared to the docked 3 RBD up conformation from PDB 7DX9.

To further define the epitope targeted by these Abs, we obtained a low-resolution cryo-EM structure of Ab16 in complex with the SARS-2 spike (**Fig. 4E-G**). The SARS-2 spike is in the “three RBD up” conformation with density for each RBD to be occupied by an Fab. Consistent with the reactivity in BLI, Ab16 appears to engage a conserved epitope that partially overlaps with the CR3022 epitope and encompasses part of the RBM (**Fig. 4E**). This latter observation will likely sterically interfere with ACE2 binding (**Fig. 4F**). Furthermore, the complex appears to show an outward rotation of the bound RBD relative to the previously characterized “three RBD up” (PDB 7DX9) conformation (**Fig. 4G**). Indeed, this was previously hypothesized to contribute to SARS-1 neutralization by CR3022 (*28*). The Ab16 binding footprint appears to overlap with previously characterized conserved epitopes targeted by antibodies with broad sarbecovirus neutralization activity: ADI-56046 and antibodies K288.2 and K398.22 isolated from rhesus macaques (*13, 36*). This region is also left unmasked in the Trimer^*hg*^ and Cocktail^*hg*^ cohort boosting immunogens (**Fig. 1**), allowing immune focusing to both conserved broadly neutralizing epitopes and the SARS-2 RBM.

Collectively, our results demonstrate immunogen design approaches that can be leveraged to enhance RBD, and more specifically, RBM-focused humoral responses. It is a strategy that maintains protective SARS-2 neutralization while also eliciting humoral responses that recognize emerging variants and coronaviruses with pandemic potential. Importantly, these design strategies are not limited to coronaviruses and are adaptable to other viruses as a general approach to elicit protective responses to conserved epitopes.

## Acknowledgments

We thank members of the Schmidt Laboratory for helpful discussions. We thank Timothy Caradonna and Catherine Jacob-Dolan for critical reading of the manuscript. We thank Dr. Jason McLellan from University of Texas, Austin for the spike plasmid. We thank Nir Hacohen and Michael Farzan for the kind gift of the ACE2 expressing 293T cells.

## Author contributions

Conceptualization, BMH, AGS; Methodology, BMH, ECL, IWW, ABB, DL, AGS; Investigation, BMH, MS, KS, ECL, JF, TK, IWW; Writing – Original Draft, BMH and AGS; Writing – Review and Editing, all authors; Funding Acquisition, ABB, DL, AGS; Supervision, ABB, DL, AGS.;

## Data and materials availability

All data is available in the main text or in the supplementary materials.

## Supplementary Materials

Materials and Methods

Figs. S1 to S14

Table S1

References (41-51)

## MATERIALS AND METHODS

### Immunogen and Coating Protein Expression and Purification

The SARS-CoV-2 (Genbank MN975262.1), SARS-CoV (Genbank ABD72970.1), WIV1-CoV (Genbank AGZ48828.1) RBDs were used as the basis for constructing these immunogens. To graft the SARS-2 RBM onto SARS-1 and WIV1 scaffolds to create the rsSARS-1 and rsWIV1 monomers, boundaries of SARS-2 residues 437 – 507 were used. All constructs were codon optimized by Integrated DNA Technologies and purchased as gblocks. Gblocks were then cloned into pVRC and sequence confirmed via Genewiz. Monomeric constructs for serum ELISA coating contained C-terminal HRV 3C-cleavable 8xHis and SBP tags. Trimeric constructs also included C-terminal HRV 3C-cleavable 8xHis tags, in addition to a previously published hyperglycosylated GCN4 tag with two engineered C-terminal cystines (*30, 31*). Dr. Jason McLellan at the University of Texas, Austin provided the spike plasmid, which contained a non-cleavable foldon trimerization domain in addition to C-terminal HRV 3C cleavable 6xHis and 2xStrep II tags. The SARS-2 Δ RBM RBD construct was generated as previously described with four additional engineered putative N-linked glycosylation sites at positions 448, 475, 494, and 501 (*30*).

Expi 293F cells (ThermoFisher) were used to express proteins. Transfections were performed with Expifectamine reagents per the manufacturer’s protocol. After 5-7 days, transfections were harvested and centrifuged for clarification. Cobalt-TALON resin (Takara) was used to perform immobilized metal affinity chromatography via the 8xHis tag. Proteins were eluted using imidazole, concentrated, and passed over a Superdex 200 Increase 10/300 GL (GE Healthcare) size exclusion column. Size exclusion chromatography was performed in PBS (Corning). For immunogens, HRV 3C protease (ThermoScientific) cleavage of affinity tags was performed prior to immunization. Cobalt-TALON resin was used for a repurification to remove the His-tagged HRV 3C protease, cleaved tag, and remaining uncleaved protein.

### Fab and IgG Expression and Purification

The variable heavy and light chain genes for each antibody were codon optimized by Integrated DNA Technologies, purchased as gblocks, and cloned into pVRC constructs which already contained the appropriate constant domains as previously described (*41, 42*). The Fab heavy chain vector contained a HRV 3C-cleavable 8xHis tag, and the IgG heavy chain vector contained HRV 3C-cleavable 8xHis and SBP tags. The same transfection and purification protocol as used for the immunogens and coating proteins was used for the Fabs and IgGs.

### Biolayer Interferometry

Biolayer interferometry (BLI) experiments were performed using a BLItz instrument (Fortebio) with FAB2G biosensors or Ni-NTA biosensors (Fortebio). All proteins were diluted in PBS. Fabs were immobilized to the biosensors, and coronavirus proteins were used as the analytes. To determine binding affinities, single-hit measurements were performed starting at 10 μM to calculate an approximate *K*_*D*_ in order to evaluate which concentrations should be used for subsequent titrations. Measurements at a minimum of three additional concentrations were performed. Vendor-supplied software was used to generate a final *K*_*D*_ estimate via a global fit model with a 1:1 binding isotherm.

### Immunizations

All immunizations were performed using female C57BL/6 mice (Jackson Laboratory) aged 6-10 weeks. Mice received 20 μg of protein adjuvanted with 50% w/v Sigma adjuvant in 100 μL of inoculum via the intraperitoneal route. Following an initial prime (day 0), boosts occurred at days 21 and 42. Serum samples were collected for characterization on day 56 from all cohorts, in addition to day 35 for the Trimer^*hg*^ and Cocktail^*hg*^ cohorts. All experiments were conducted with institutional IACUC approval (MGH protocol 2014N000252).

### Serum ELISAs

Serum ELISAs were executed using 96-well, clear, flat-bottom, high bind microplates (Corning). These plates were coated with 100 μL of protein, which were adjusted to a concentration of 5 μg/mL (in PBS). Plates were incubated overnight at 4°C. After incubation, plates had their coating solution removed and were blocked using 1% BSA in PBS with 1% Tween. This was done for 60 minutes at room temperature. This blocking solution was removed, and sera was diluted 40-fold in PBS. A 5-fold serial dilution was then performed. CR3022 IgG, similarly serially diluted (5-fold) from a 5 μg/mL starting concentration, was used as a positive control. 40 μL of primary antibody solution was used per well. Following this, samples were incubated for 90 minutes at room temperature. Plates were washed three times using PBS-Tween. 150 μL of HRP-conjugated rabbit anti-mouse IgG antibody, sourced commercially from Abcam (at a 1:20,000 dilution in PBS), was used for the secondary incubation. Secondary incubation was performed for one hour, similarly at room temperature. Plates were subsequently washed three times using PBS-Tween. 1xABTS development solution (ThermoFisher) was used according to the manufacturer’s protocol. Development was abrogated after 30 minutes using a 1% SDS solution, and plates were read using a SectraMaxiD3 plate reader (Molecular Devices) for absorbance at 405 nm.

### Competition ELISAs

A similar protocol to the serum ELISAs was used for the competition ELISAs. For the primary incubation, 40 μL of the relevant IgG at 1 μM was used at room temperature for 60 minutes. Mouse sera were then spiked in such that the final concentration of sera fell within the linear range for the serum ELISA titration curve for the respective coating antigen, and an additional 60 minutes of room temperature incubation occurred. After removing the primary solution, plates were washed three times with PBS-Tween. Secondary incubation consisted of HRP-conjugated goat anti-mouse IgG, human/bovine/horse SP ads antibody (Southern Biotech) at a concentration of 1:4000. The remaining ELISA procedure (secondary incubation, washing, developing) occurred as described for the serum ELISAs. Percent binding loss was calculated relative to a no IgG control. Negative percent binding loss values were set to zero for the purpose of visualizations.

### ACE2 Cell Binding Assay

ACE2 expressing 293T cells (*43*) (a kind gift from Nir Hacohen and Michael Farzan) were harvested. A wash was performed using PBS supplemented with 2% FBS. 200,000 cells were allocated to each labelling condition. Primary incubation occurred using 100 μL of 1 μM antigen in PBS on ice for 60 minutes. Two washes were performed with PBS supplemented with 2% FBS. Secondary incubation was performed using 50 μL of 1:200 streptavidin-PE (Invitrogen) on ice for 30 mins. Two washes were performed with PBS supplemented with 2% FBS, and then cells were resuspended in 100 μL of PBS supplemented with 2% FBS. A Stratedigm S1000Exi Flow Cytometer was used to perform flow cytometry. FlowJo (version 10) was used to analyze FCS files.

### Pseudovirus Neutralization Assay

Serum neutralization against SARS-CoV-2, SARS-CoV, WIV1-CoV, RaTG13, and SHC014 was assayed using pseudotyped lentiviral particles expressing spike proteins described previously (*32*). Transient transfection of 293T cells was used to generate lentiviral particles. Viral supernatant titers were measured using flow cytometry of 293T-ACE2 cells (*43*) and utilizing the HIV-1 p24^CA^ antigen capture assay (Leidos Biomedical Research, Inc.). 384-well plates (Grenier) were used to perform assays on a Tecan Fluent Automated Workstation. For mouse sera, samples underwent primary dilutions of 1:3 or 1:9 followed by serial 3-fold dilutions. 20 μL each of sera and pseudovirus (125 infectious units) were loaded into each well. Plates were then incubated for 1 hour at room temperature. Following incubation, 10,000 293T-ACE2 cells (*43*) in 20 μL of media containing 15 μg/mL polybrene was introduced to each well. The plates were then further incubated at 37°C for 60-72 hours.

Cells were lysed using assay buffers described previously (*44*). Luciferase expression was quantified using a Spectramax L luminometer (Molecular Devices). Neutralization percentage for each concentration of serum was calculated by deducting background luminescence from cells-only sample wells and subsequently dividing by the luminescence of wells containing both virus and cells. Nonlinear regressions were fitted to the data using GraphPad Prism (version 9), allowing IC50 values to be calculated via the interpolated 50% inhibitory concentration. IC50 values were calculated with a neutralization values greater than or equal to 80% at maximum serum concentration for each sample. NT50 values were then calculated using the reciprocal of IC50 values. Serum neutralization potency values were calculated by dividing the NT50 against a particular pseudovirus by the endpoint titer against the respective RBD. For samples with NT50 values below the limit of detection, the lowest limit of detection across all neutralization assays was used as the NT50 value to calculate neutralization potency. This prevents a higher limit of detection from skewing neutralization potency results. Endpoint titers were normalized relative to a CR3022 IgG control, which was run in every serum ELISA. ELISA titers that were too low to calculate an endpoint titer were set to 40, which was the starting point for the serum dilutions.

In comparing NT50 values for the various cohorts across the wildtype and variant pseudoviruses, the lowest limit of detection across all neutralization assays performed for a given cohort was used for any NT50 values that fell below the limit of detection. This prevents a higher limit of detection in some assays from skewing the comparison results.

### Flow Cytometry

Single cell suspensions were generated from mouse spleens following isolation via straining through a 70 μm cell strainer. Treatment with ACK lysis buffer was performed to remove red blood cells, and cells were washed with PBS. Aqua Live/Dead amine-reactive dye (0.025 mg/mL) was first used to stain single cell suspensions. The following B and T cell staining panel of mouse-specific antibodies was then applied: CD3-BV786 (BioLegend), CD19-BV421 (BioLegend), IgM-BV605 (BioLegend), IgG-PerCP/Cy5.5 (BioLegend). Staining was performed using a previously described staining approach (*45, 46*).

SBP-tagged coronavirus proteins were labelled using streptavidin-conjugated flurophores as previously described (*47*). Briefly, a final conjugated probe concentration of 0.1 μg/mL was achieved following the addition of streptavidin conjugates to achieve a final molar ratio of probe to streptavidin valency of 1:1. This addition was performed in 5 increments with 20 minutes of incubation at 4°C with rotation in between. The coronavirus protein panel consisted of the following flurorescent probes: SARS-CoV-2 RBD-APC/Cy7 (streptavidin-APC/Cy7 from BioLegend), WIV1 RBD-BV650 (streptavidin-BV650 from BioLegend), SARS-CoV-2 spike-StreptTactin PE (StrepTactin PE from IBA Lifesciences), and SARS-CoV-2 spike-StreptTactin APC (StrepTactin APC from IBA Lifesciences).

A BD FACSAria Fusion cytometer (BD Biosciences) was used to perform flow cytometry. FlowJo (version 10) was used to analyze the resultant FCS files. Sorted cells were IgG+ B cells that were double-positive for SARS-CoV-2 spike and positive for the SARS-CoV-2 RBD.

### B Cell Receptor Sequencing

Cells were sorted into 96-well plates containing 4 μL of lysis buffer, consisting of 0.5X PBS, 10 mM DTT, and 4 units of RNaseOUT (ThermoFisher). Following sorting, plates were spun down at 3000g for 1 minute and stored at -80°C. Plates were later thawed and a reverse transcriptase reaction was performed using the SuperScript IV VILO MasterMix (ThermoFisher) in a total volume of 20 μL according to the manufacturer’s recommendations. Two rounds of PCR were then performed using previously published primers (*48, 49*). Variable heavy and light chains were then sequenced via Sanger sequencing (Genewiz).

IMGT High V-Quest was used to analyze variable heavy and light chain sequences, and IgBlast was used to identify clonal lineages. Data were plotted using Python.

### Cryo-EM Grid Preparation and Image Recording

Complexes of SARS-CoV-2 spike (6P) with Ab16 Fab were formed by combining spike at 0.7 mg/mL with Fab at 0.6 mg/mL (three-fold excess of binding sites) in a buffer composed of 10 mM Tris pH 7.5 with 150 mM NaCl. Spike·Fab complexes were incubated for 30 minutes on ice before application to thick C-flat 1.2-1.3 400 Cu mesh grids (Protochips). Grids were glow discharged (PELCO easiGlow) for 30 seconds at 15 mA and prepared with a Gatan Cryoplunge 3 by applying 3.8 uL of sample and blotting for 4.0 seconds in the chamber maintained at a humidity between 88% and 92%. Images for Spike complexes with Ab16 were recorded on a Talos Arctica microscope operated at 200 keV with a Gatan K3 direct electron detector. Automated image acquisition was performed with Serial EM (*50*).

### Cryo-EM Image Analysis and 3D Reconstruction and Model Fitting

Image analysis for was carried out in RELION as previously. Briefly, particles were extracted from motion-corrected micrographs and subjected to 2D classification, initial 3D model generation, 3D classification, and 3D refinement. 2D class averages are shown in **Fig. S14**. Ab16 was C3 symmetric. CTF refinement was performed to correct beam tilt, trefoil, anisotropic magnification, and per particle defocus in RELION (*51*). Bayesian polishing was also performed in RELION leading to a 6.6 Å reconstruction following 3D refinement. The final 3D refined map was sharpened with a B-factor of −297.5 Å^2^ resulting in a 5.5 Å resolution map as determined by the Fourier shell correlation (0.143 cutoff). Heavy and light chains of PDB entries 4L5F and 4HC1 were aligned and extracted to make an initial model for the Fab. Spike with 3 RBD in the “up” conformation (PDB 7DX9) and model of Ab16 Fab were docked into the cryoEM map using Chimera.

### Statistical Analysis

Curve fitting and statistical analyses were performed with GraphPad Prism (version 9). Non-parametric statistics were used throughout. To compare multiple populations, the Kruskal-Wallis non-parametric ANOVA was used with post hoc analysis using Dunn’s test for multiple comparisons. The Mann-Whitney U test was used to compare two populations without consideration for paired samples. The ratio-paired t-test was used to compare two populations with consideration for paired samples and evidence of normality. P values in ANOVA analyses were corrected for multiple comparisons. A p value < 0.05 was considered significant.

