## Supplemental Materials for "Rationally designed immunogens enable immune focusing to the SARS-CoV-2 receptor binding motif"

**This PDF file includes:**

Materials and Methods
Figs. S1 to S14
Table S1

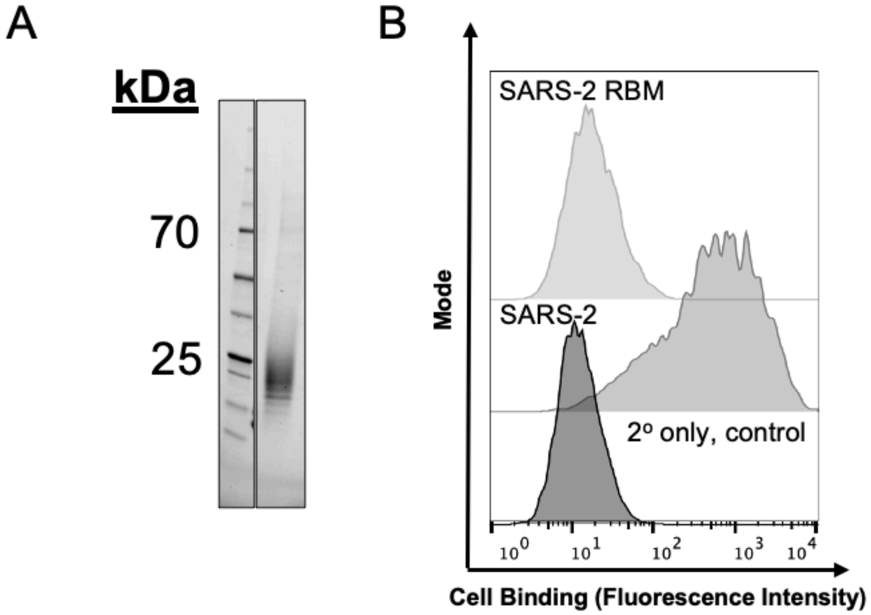


**Figure S1. SARS-CoV-2 receptor binding motif expression.** (**A**) SDS-Page gel analysis of the SARS-2 receptor binding motif (RBM) construct with HRV 3C-cleavable 8xHis and streptavidin binding peptide (SBP) tags. (**B**) ACE2 cell binding assay results for the SARS-2 RBM construct at 1 μM.

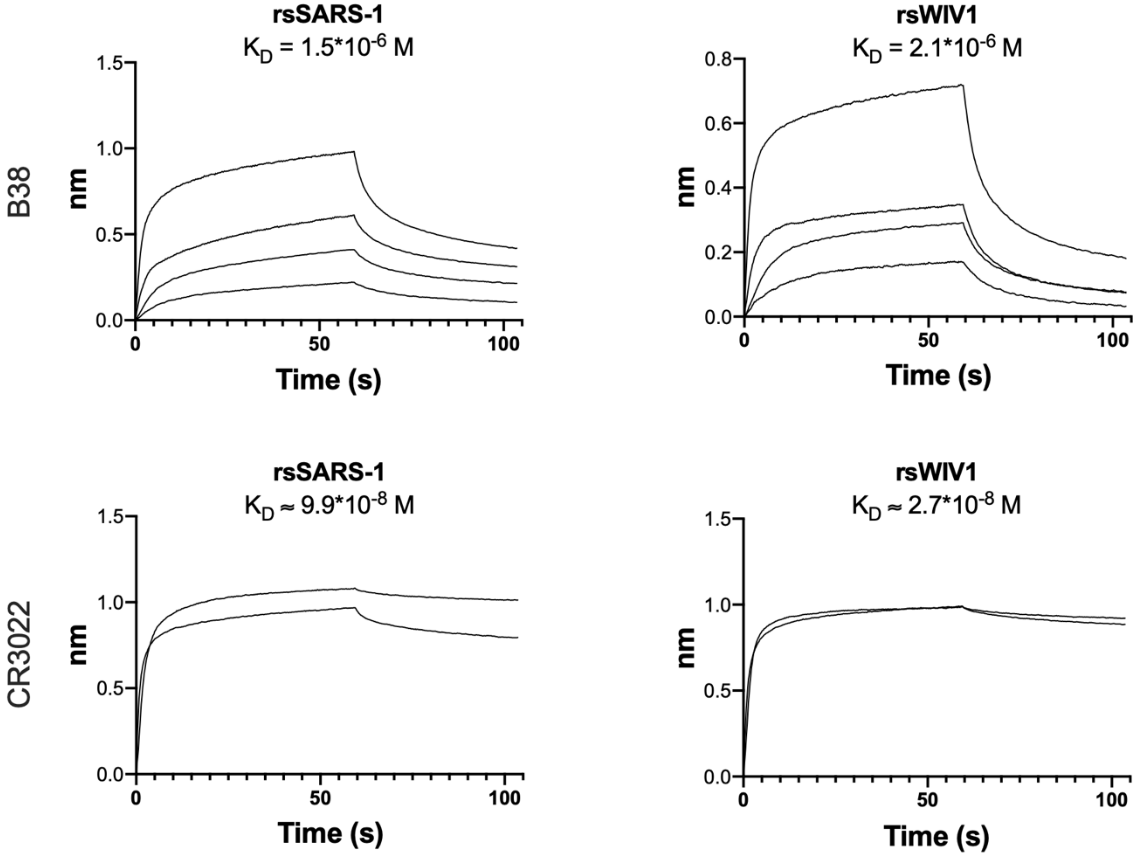


**Figure S2. Resurfaced SARS-1 and WIV1 BLI.** After grafting the SARS-2 RBM onto the SARS-1 and WIV1 RBDs, conformationally specific Fabs B38 and CR3022 were used to confirm that these epitopes remained intact with comparable affinity to wildtype (9.7*10$^{-7}$ M for B38 Fab and SARS-2 RBD; 2.7*10$^{-8}$ M for CR3022 Fab and SARS-1 RBD; 3.7*10$^{-8}$ M for CR3022 Fab and WIV1 RBD). FAB2G sensors were used with immobilized fabs. rsSARS-1 and rsWIV1 were the analytes. Titrations with B38 Fab were performed at 10 µM, 5 µM, 1 µM, and 0.5 µM. Titrations with CR3022 Fab were performed at 10 µM and 5 µM. Vendor-supplied software was used to generate an apparent $K_D$, or an approximate $K_D$ in the case of titrations with two runs.

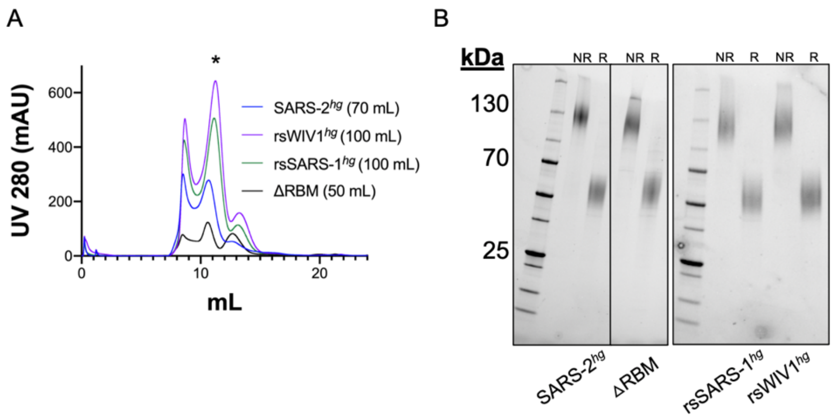


C

| Trimer | mg per 1L Expi293 |
| --- | --- |
| SARS-2$^{hg}$ | 21 |
| rsSARS-1$^{hg}$ | 25 |
| rsWIV1$^{hg}$ | 33 |
| ΔRBM | 11 |

**Figure S3. Trimer expression and purification.** (**A**) Representative size exclusion chromatography traces for trimeric constructs. The trimer peak is marked with "*", and fractions from this peak were pooled for HRV 3C cleavage and use as immunogens. Quantity of Expi293 transfection is in parentheses next to each label. (**B**) SDA-Page gel analysis of purified and cleaved trimeric constructs under non-reducing (NR) and reducing (R) conditions, showing the dissociation of the disulfide bond in the hyperglycosylated GCN4 tag (hgGCN4$^{cys}$) under reducing conditions. (**C**) Protein yields for purified trimeric constructs in Expi293 cells.

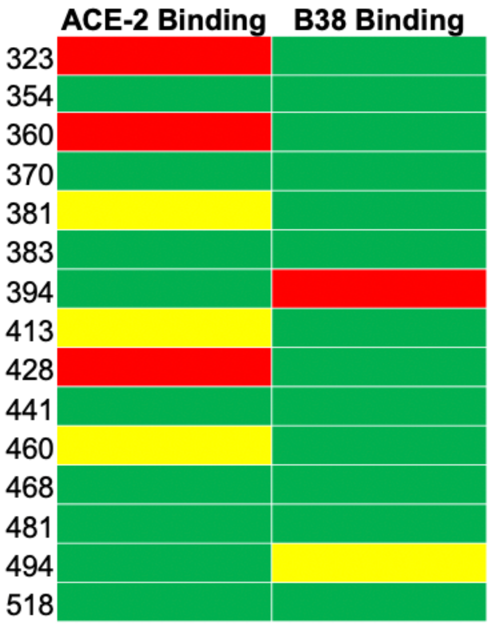


**Figure S4. Single glycan biochemical validation.** Candidate glycans were tested individually in the context of the SARS-CoV-2 RBD. All glycans were designed to mask epitopes outside the RBM. Therefore, biochemical validation was performed by assessing binding to the RBM-directed conformationally specific antibody B38 via single-hit BLI using FAB2G sensors with the RBD of interest as the analyte at 10 μM. Binding to ACE2 was also assessed via an ACE2 cell binding assay with antigen concentrations at 1 μM. Binding was binned subjectively into three categories: minimal (red), substantially reduced (yellow), and roughly intact (green).

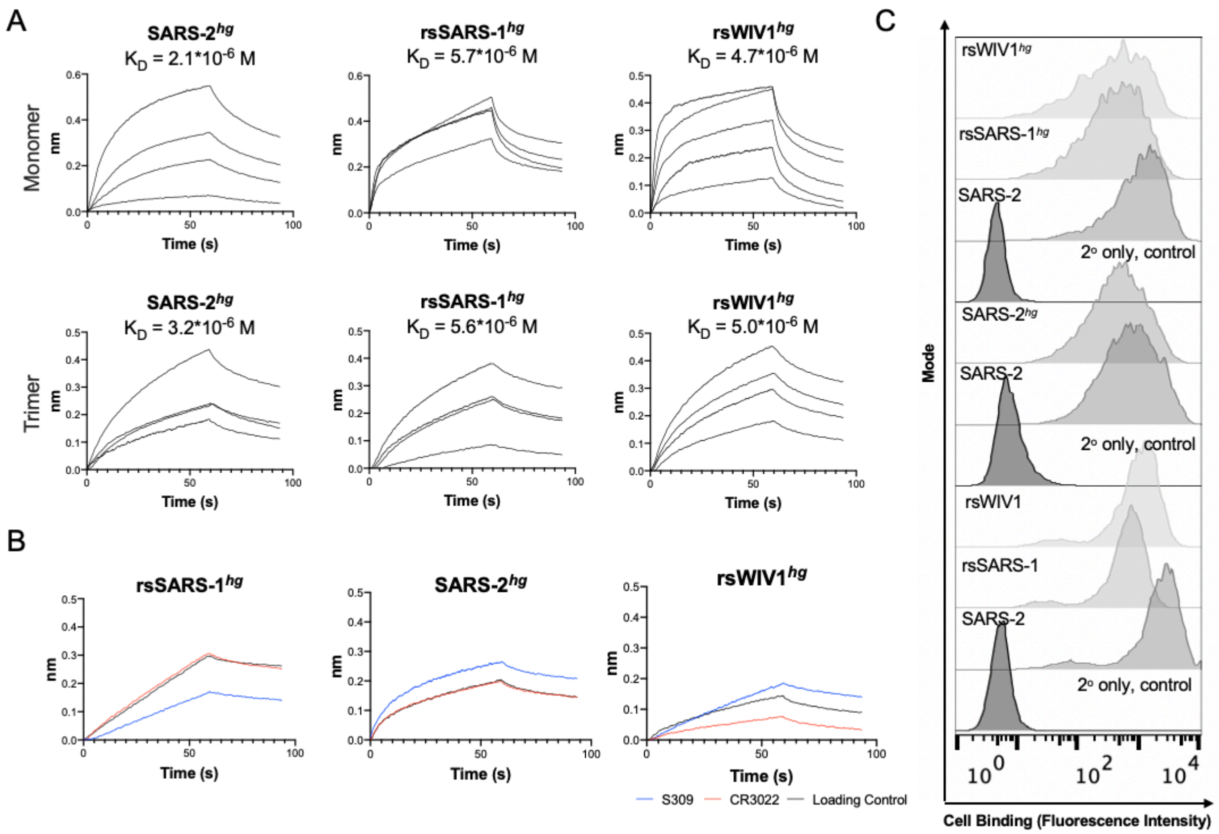


**Figure S5. Biochemical validation of hyperglycosylated RBDs.** (**A**) Conformationally specific Fab B38 was used to assess continued accessibility of the SARS-2 RBM in both monomeric and trimeric hyperglycosylated constructs. FAB2G sensors were used with immobilized fabs; hyperglycosylated coronavirus proteins were the analytes. Titrations with B38 Fab were performed at 7.5 μM, 5 μM, 2.5 μM, and 1 μM (rsSARS-1$^{hg}$ trimer, rsWIV1$^{hg}$ trimer); 5 μM, 2.5 μM, 1 μM, and 0.5 μM (SARS-2$^{hg}$ trimer, SARS-2$^{hg}$ monomer); 10 μM, 7.5 μM, 5 μM, 3.75 μM, and 2.5 μM (rsWIV1$^{hg}$ monomer); 10 μM, 7.5 μM, 5 μM, and 2.5 μM (rsSARS-1$^{hg}$ monomer). Vendor-supplied software was used to generate an apparent $K_D$. (**B**) BLI with conformationally specific Fabs CR3022 and S309 at 10 μM was compared to loading controls for each of the hyperglycosylated monomers to confirm a lack of binding. (**C**) ACE2 cell binding assay results for resurfaced and hyperglycosylated constructs at 1 μM.

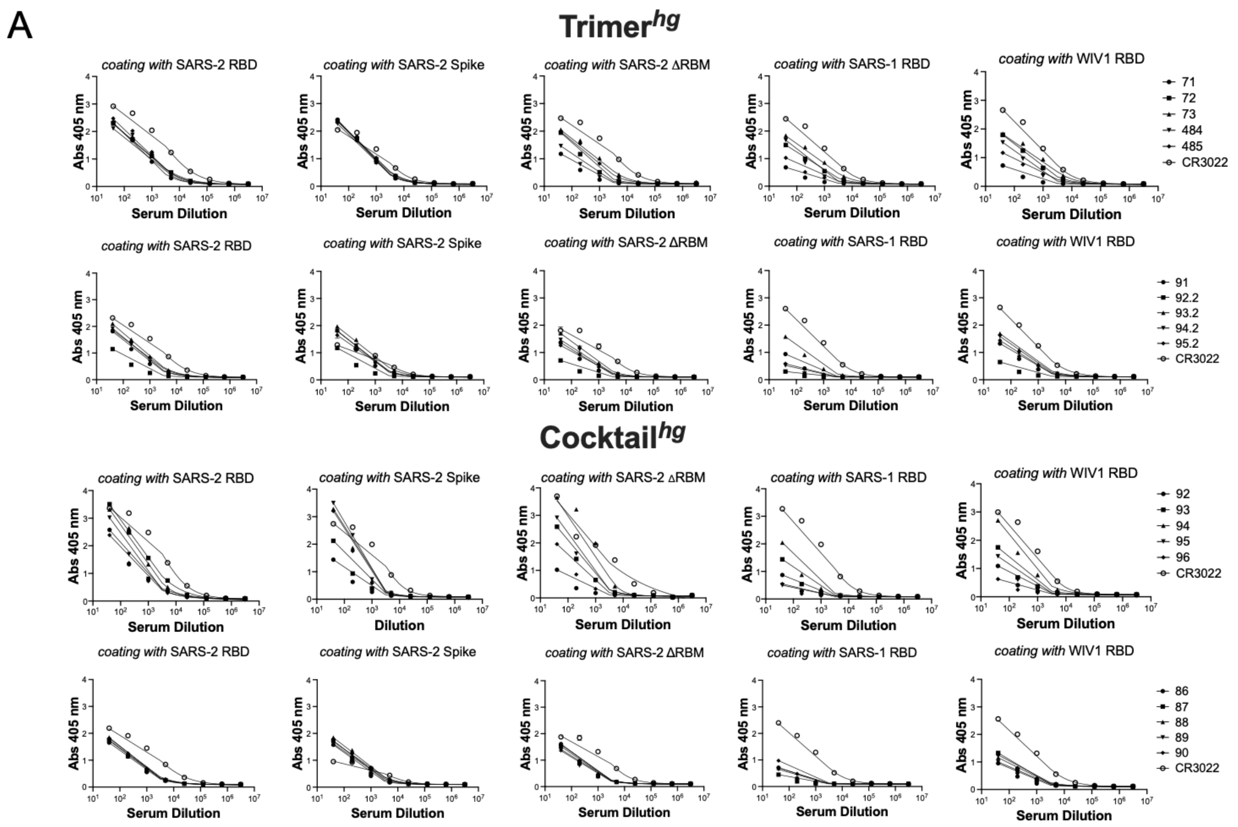
A
Trimer^hg
coating with SARS-2 RBD
coating with SARS-2 Spike
coating with SARS-2 ΔRBM
coating with SARS-1 RBD
coating with WIV1 RBD
Abs 405 nm
Serum Dilution
71
72
73
484
485
CR3022
91
92.2
93.2
94.2
95.2
CR3022
Cocktail^hg
Dilution
92
93
94
95
96
CR3022
86
87
88
89
90
CR3022

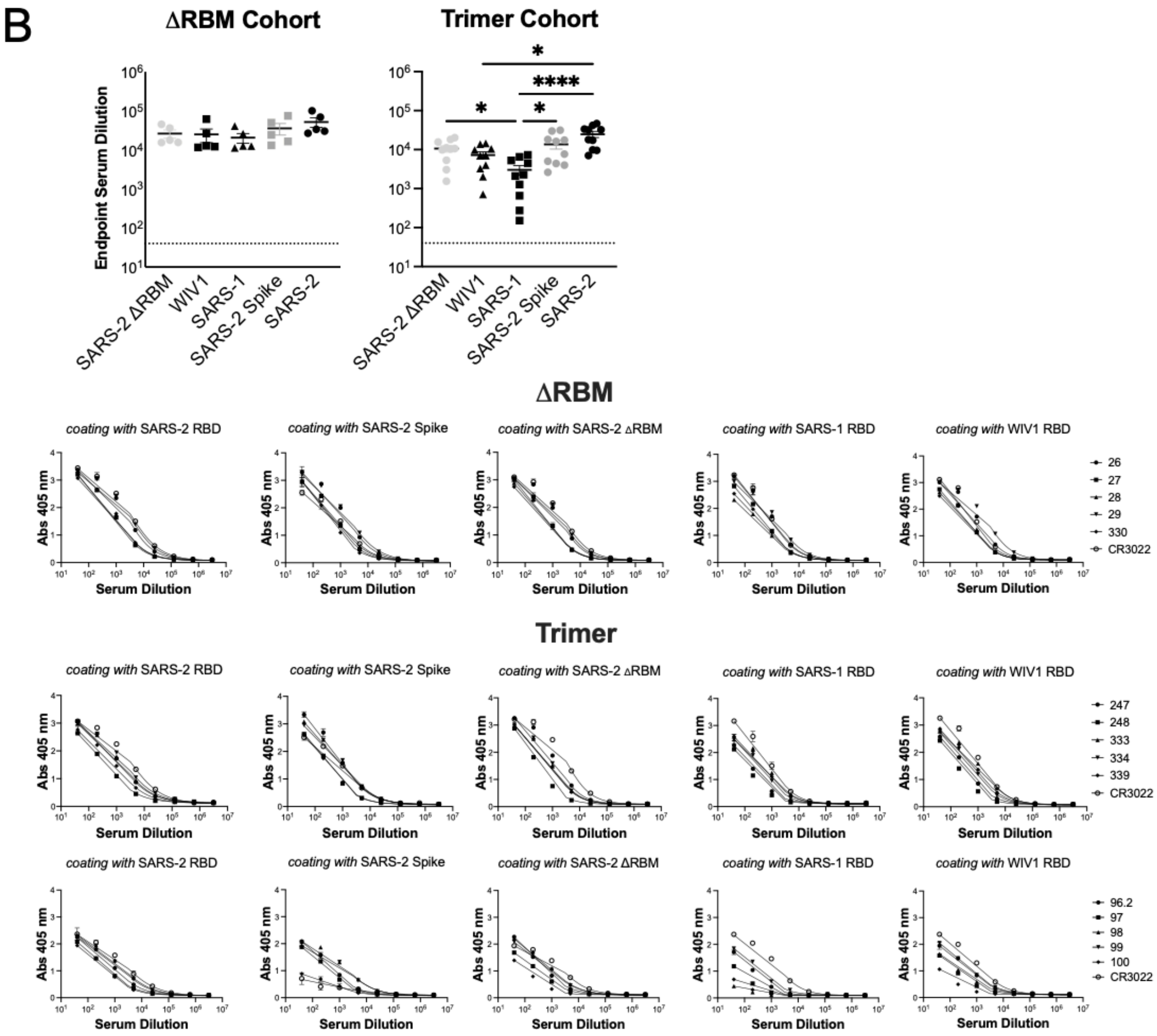

B
ΔRBM Cohort
Trimer Cohort
Endpoint Serum Dilution
SARS-2 ΔRBM
WIV1
SARS-1
SARS-2 Spike
SARS-2
*
****
ΔRBM
coating with SARS-2 RBD
coating with SARS-2 Spike
coating with SARS-2 ΔRBM
coating with SARS-1 RBD
coating with WIV1 RBD
Abs 405 nm
Serum Dilution
26
27
28
29
330
CR3022
Trimer
247
248
333
334
339
CR3022
96.2
97
98
99
100
CR3022

C

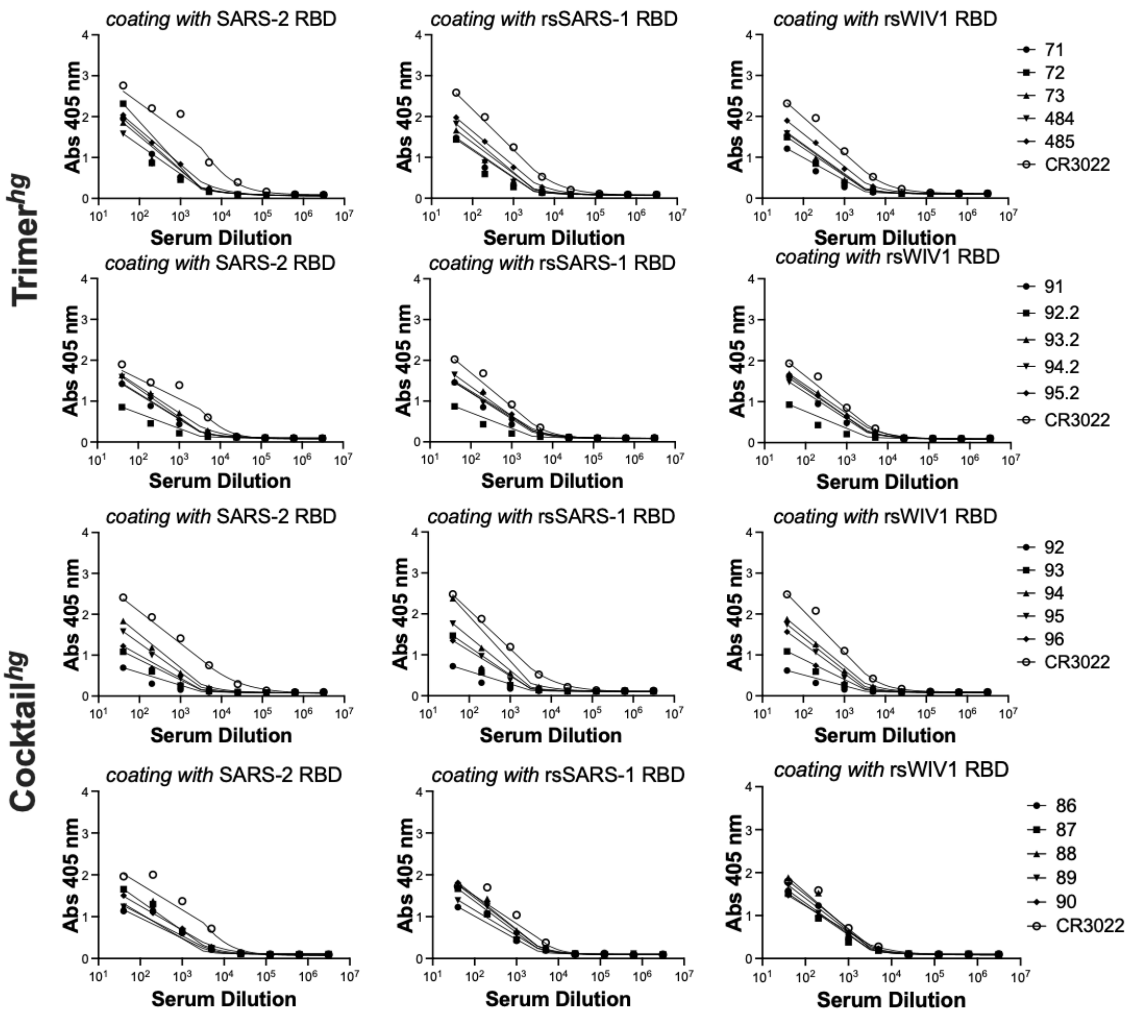
Trimer^hg
Cocktail^hg
coating with SARS-2 RBD
coating with rsSARS-1 RBD
coating with rsWIV1 RBD
Abs 405 nm
Serum Dilution
71
72
73
484
485
CR3022
91
92.2
93.2
94.2
95.2
CR3022
92
93
94
95
96
CR3022
86
87
88
89
90
CR3022

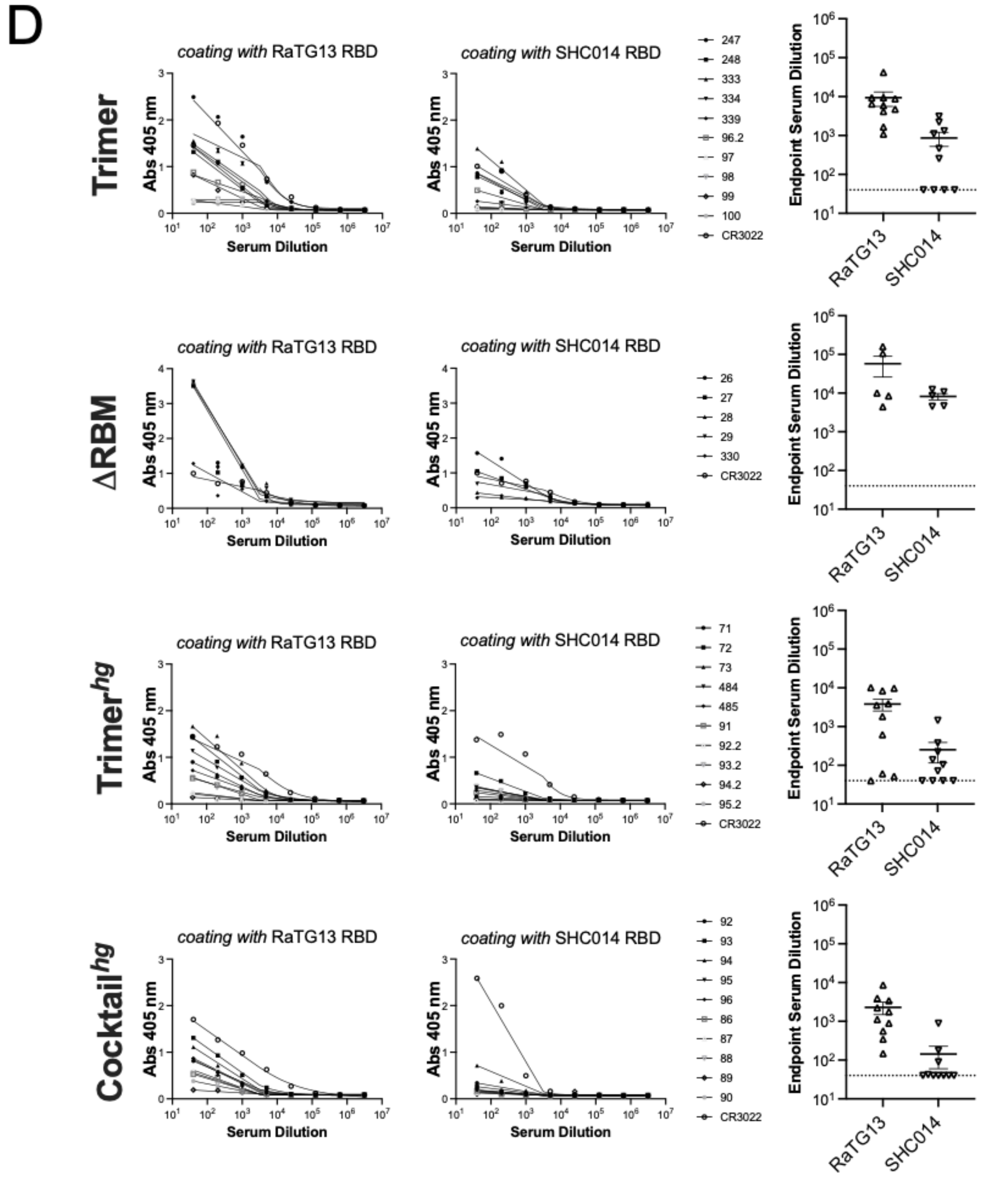


**Figure S6. Serum ELISAs with coronavirus proteins.** Serum ELISAs were performed using day 56 serum samples to assess binding to coronavirus proteins. These assays were performed on (**A**) The Trimer$^{hg}$ and Cocktail$^{hg}$ cohorts and (**B**) the ΔRBM and Trimer cohorts. Endpoint titers were calculated based on curves fit using a sigmoidal model. (**C**) Binding was also assayed against the rsSARS-1 and rsWIV1 RBDs using sera from the Trimer$^{hg}$ and Cocktail$^{hg}$ cohorts. (**D**) For all cohorts, day 56 serum samples were used in ELISAs to assess binding to RaTG13 and SHC014 RBDs.

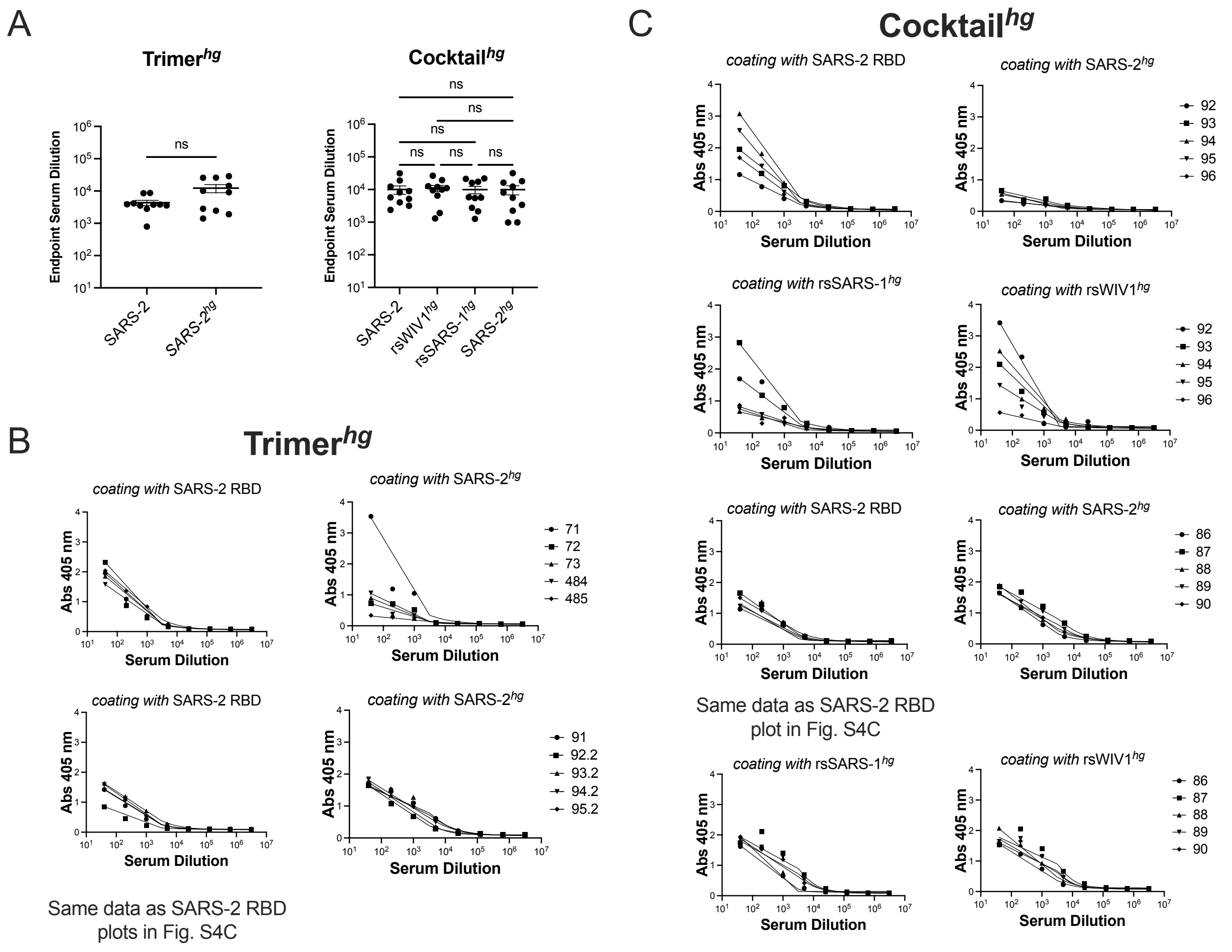


**Figure S7. Serum ELISAs with hyperglycosylated immunogens.** Serum ELISAs were performed against the relevant hyperglycosylated immunogens for the Trimer$^{hg}$ and Cocktail$^{hg}$ cohorts. (**A**) Endpoint titers against all hyperglycosylated immunogens do not vary significantly from titers against wildtype SARS-2 RBD. Endpoint titers were calculated using curves fit with a sigmoidal model to data from the (**B**) Trimer$^{hg}$ and (**C**) Cocktail$^{hg}$ cohorts. There was no statistically significant difference in binding within the Trimer$^{hg}$ or Cocktail$^{hg}$ cohorts across these coating antigens, as determined by the Mann-Whitney U test and the Kruskal-Wallis test, respectively.

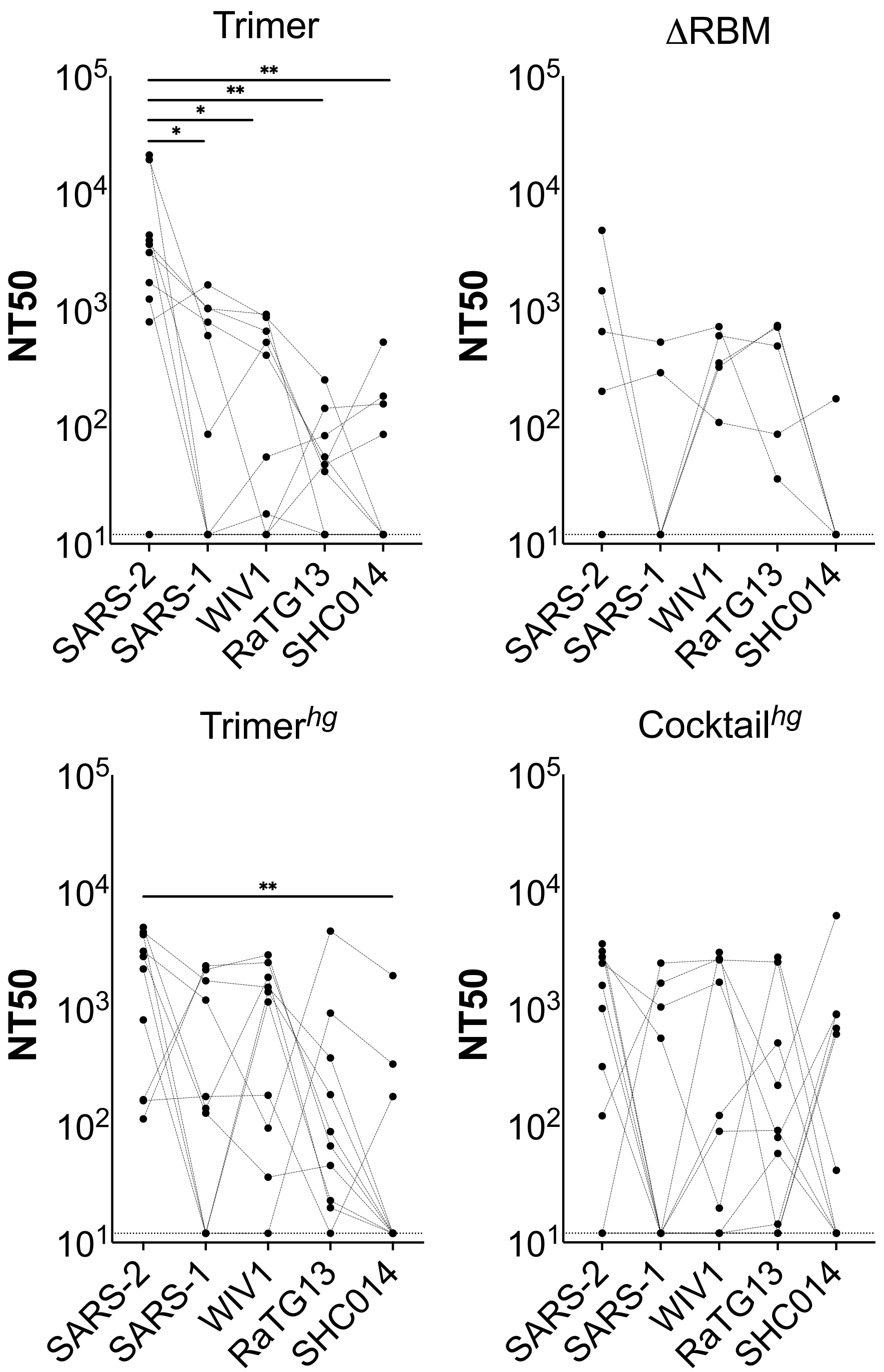


**Figure S8. Serum neutralization titers organized by cohort.** Pseudovirus neutralization assays were used to calculate NT50 values for SARS-2, SARS-1, WIV1, RaTG13, and SHC014 from all cohorts. All NT50s are from day 56 sera. Statistical significance was determined using the Friedman test to account for sample pairing with post-hoc analysis using Dunn's test corrected for multiple comparisons (* = p < 0.05, ** = p < 0.01).

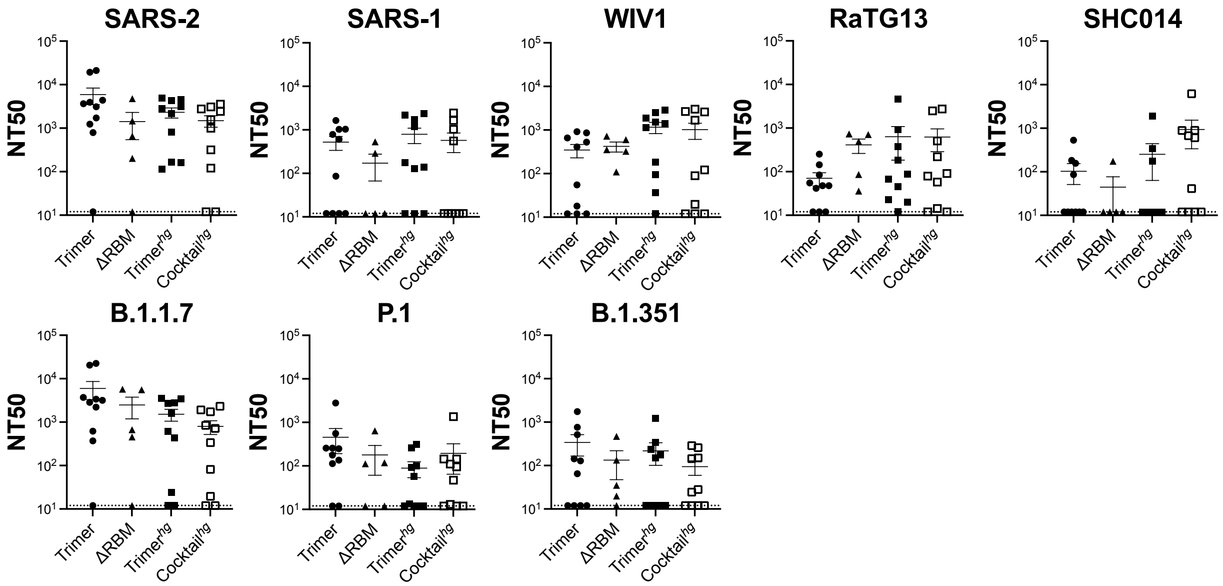


**Figure S9. Serum neutralization titers organized by pseudovirus.** Pseudovirus neutralization assays were used to calculate NT50 values for SARS-2, SARS-1, WIV1, RaTG13, and SHC014 from all cohorts. All NT50s are from day 56 sera. Statistical significance was determined using the Kruskal-Wallis test with post-hoc analysis using Dunn's test corrected for multiple comparisons; no differences are statistically significant.

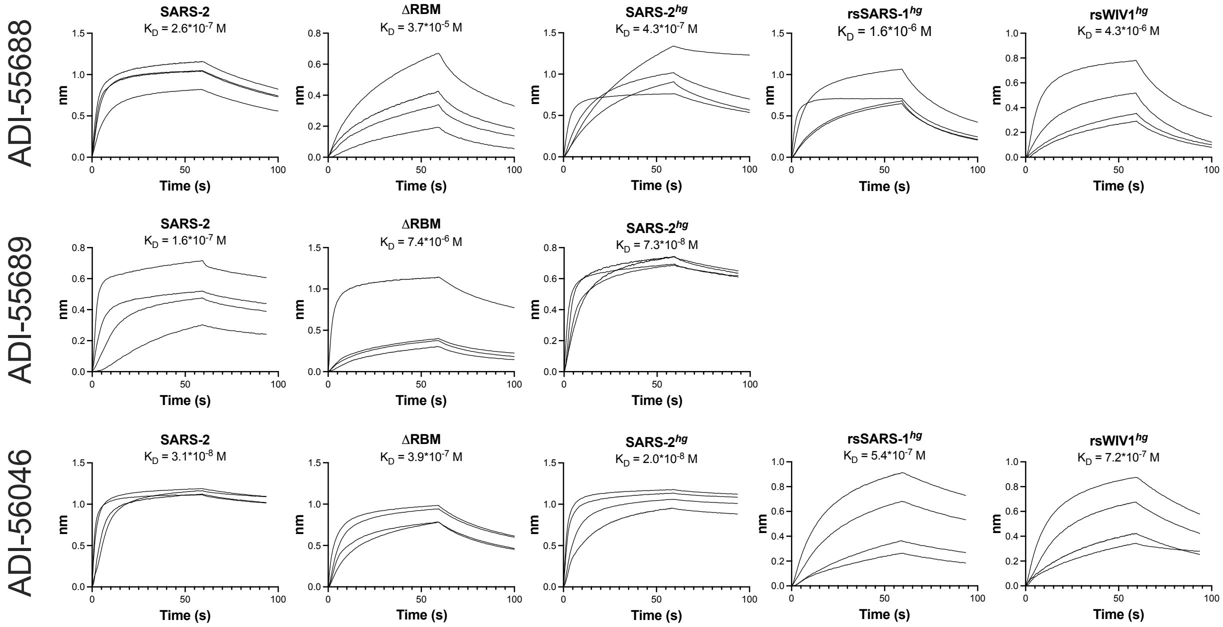


**Figure S10. BLI with antibodies to a conserved RBM epitope.** Conformationally specific Fabs ADI-55688, ADI-55689, and ADI-56046, which target a conserved RBM epitope, were used to assess binding to the ΔRBM, SARS-2$^{hg}$, rsSARS-1$^{hg}$, and rsWIV1$^{hg}$ monomers via BLI. FAB2G sensors were used with immobilized fabs; coronavirus proteins were the analytes. Titrations were performed at 10 μM, 5 μM, 2.5 μM, and 1.25 μM (SARS-2 RBD and SARS-2$^{hg}$ with ADI-55689, SARS-2 RBD and SARS-2$^{hg}$ with ADI-56046, all titrations with ΔRBM); 3 μM, 1.5 μM, 0.75 μM, and 0.375 μM (rsSARS-1$^{hg}$ and rsWIV1$^{hg}$ with ADI-56046); 10 μM, 5 μM, 2.5 μM, and 1 μM (all titrations with ADI-55688 except ΔRBM). Minimal binding was detected to ADI-55689 Fab with rsSARS-1$^{hg}$ and rsWIV1$^{hg}$. Vendor-supplied software was used to generate an apparent $K_D$.

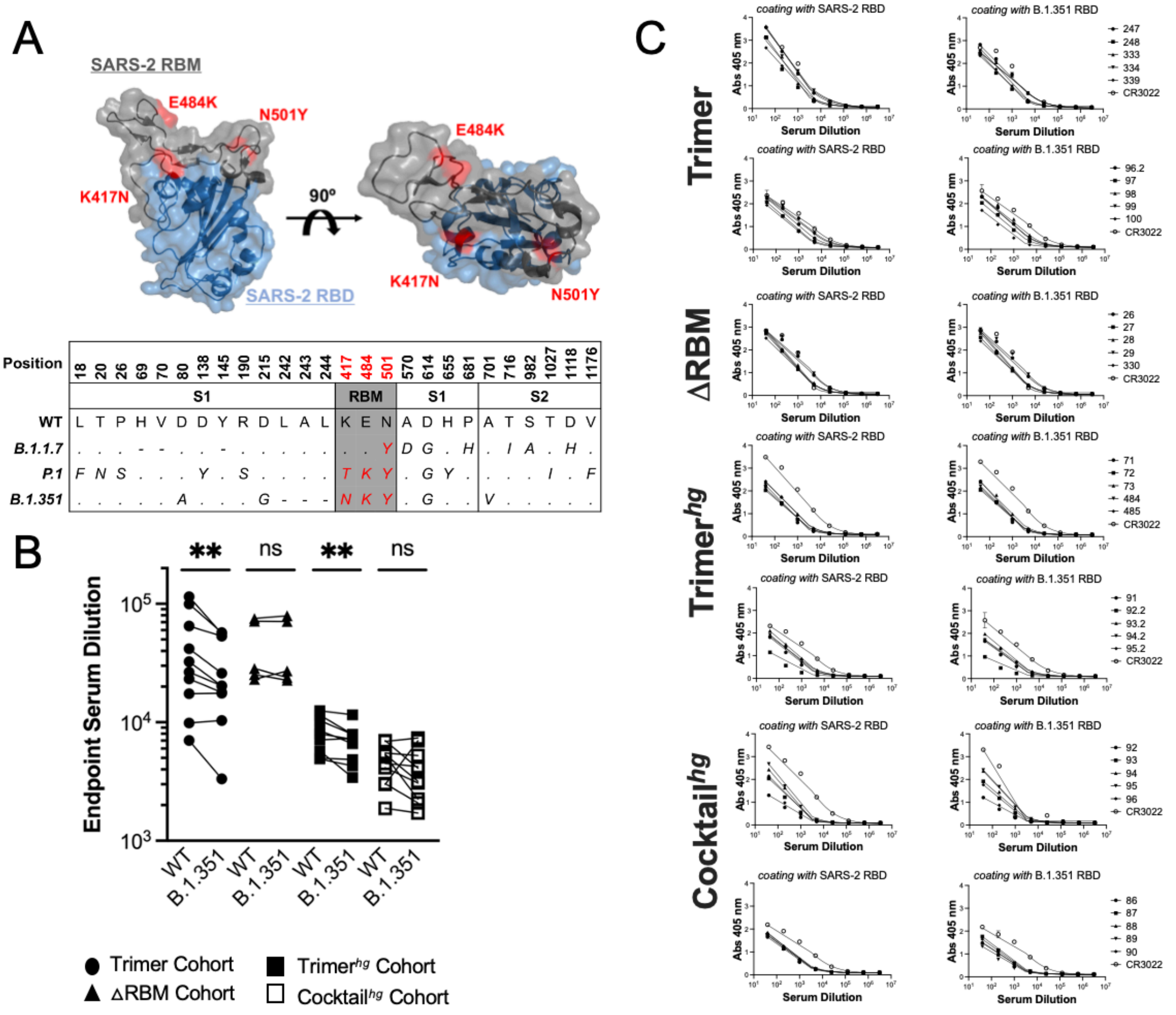


**Fig. S11. Binding of SARS-2 variants. (A)** Structural depiction of SARS-2 variant RBD mutations for 501Y.V2 (red), as well as ACE2 contact residues (cyan). (PDB: 6M0J) Sequences depict all spike mutations across select variants. **(B)** Day 56 serum was assayed in ELISA against SARS-2 RBD (WT) and SARS-2 RBD with K417N, E484K, and N501Y mutations (B.1.351). Statistical significance was determined using the Wilcoxon sign rank test (* = $p < 0.05$, ** = $p < 0.01$; ns = not significant). **(C)** Endpoint titers were calculated based on curves fit to ELISA data using a sigmoidal model.

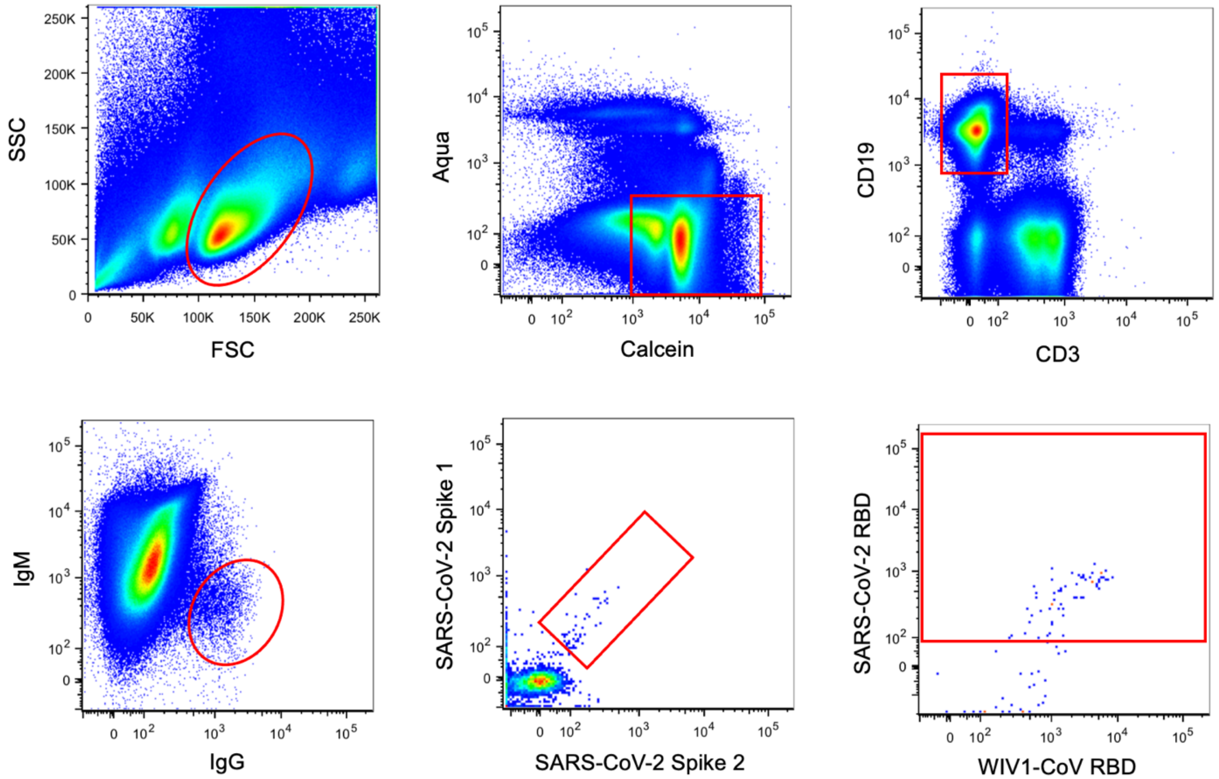


**Figure S12. Sorting scheme.** Gating scheme for isolating IgG+ B cells that are SARS-CoV-2 spike double-positive and SARS-CoV-2 RBD positive.

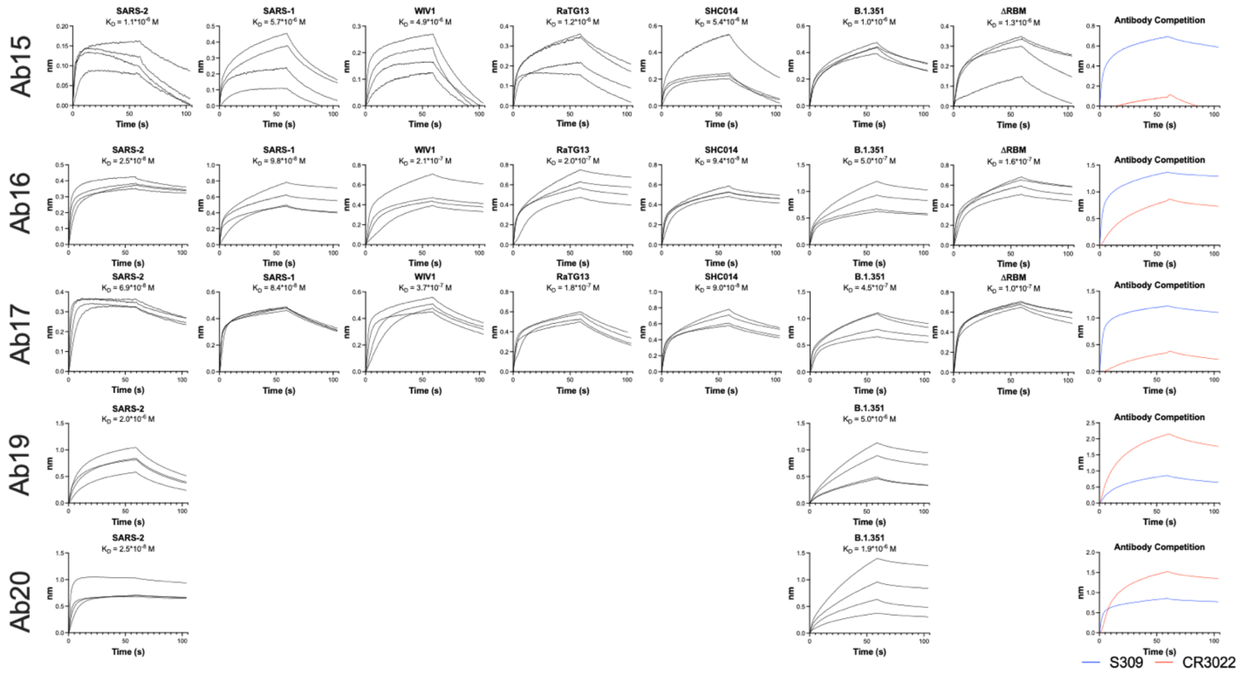


**Fig. S13. BLI with Fabs representative of lineages expanded in RBM-focusing cohorts.** BLI was performed using Fabs representative of lineages expanded in RBM-focusing cohorts. Ni-NTA biosensors were used with Fabs bound to the sensor using the 8xHis tag and RBDs in solution as the analyte. Titrations were performed at 10 μM, 5 μM, 2.5 μM, and 1 μM. Vendor-supplied software was used to generate an apparent $K_D$.

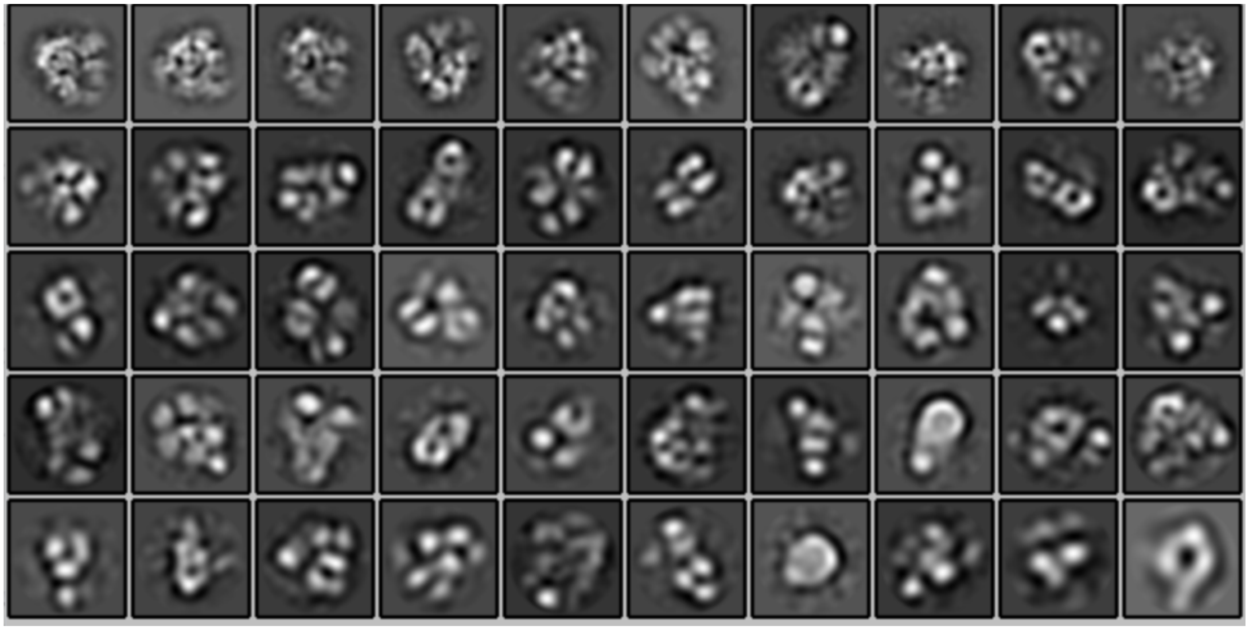

**Fig. S14. 2D class averages of the SARS-CoV-2 Spike:Ab16 complex.** Selected 2D class averages generated for the SARS-CoV-2 spike in complex with Ab16 Fab.

| Antibody | VH Sequence | VL Sequence |
|---|---|---|
| Ab15 | GAGGTCCAGCTGCAGCAGTCTGGACCTGAGCTGGTGAAGCCTGGGGCTTCAGT<br>GAAGATATCCTGCAAGGCTTCTGGTTACTCATTCACTGGCTACTACATGAACTGG<br>GTGAAGCAAAGTCCTGAAAAGAGCCTTGAGTGGATTGGAGAGATTAATCCTAACTT<br>TGGTGGTACTACCTACAACCAGAAGTTCAAGGCCAAGGCCACATTGACTGTAGAC<br>AAATCCTCCAGCACAGCCTACATGCAGCTCAAGAGCCTGACATCTGAGGACTCTG<br>CAGTCTATTACTGTGCAAGATACTATGGTAACCTCTATGCTATGGACTACTGGGGT<br>CAAGGAACCTCAGTCACCGTCTCC | GACATCCTGATGACCCAGTCTCCATCCTCCATGTCTGTATCTCTGGGAGACACAG<br>TCAGCATCACTTGCCATGCAAGTCAGGGCATTAGCAGTAATATAGGGTGGTTGCA<br>GCAGAAACCAGGGAAATCATTTAAGGGCCTGATCTATCATGGAACCAACTTGGAA<br>GATGGAGTTCCATCAAGGTTCAGTGGCAGTGGATCTGGAGCAGCTTATTCTCTCA<br>CCATCAGCAGCCTGGAATCTGAAGATTTTGCAGACTATTACTGTGTACAGTATACT<br>CATTTTCCGTACACGTTCGGAGGGGGGACCAAGCTGGAAATAAAA |
| Ab16 | GAGGTCCAGCTGCAGCAGTCTGGACCTGAGCTGGTGAAGCCTGGGGCTTCAGT<br>GAAGATATCCTGCAAGGCTTCTGGTTACTCATTTAATAACTACTACATGAACTGGG<br>TGAAGCAGAGTCCTGAAAAGAGCCTTGAGTGGATTGGAGAGATTAATCCTAACTC<br>TGGTTATACTTCCTACAACCAGAAGTTCAGGGCCAAGGCCACATTGACTGTAGACA<br>AATCCTCCACCACAGCCTACATGCAGCTCAAGAGCCTGACATCTGAGGACTCTGC<br>GGTCTATTACTGTGCAAGATACTTTGGTAACCTCTTTGCTATGGACTTCTGGGGT<br>CAAGGAACCTCAGTCACCGTCTCC | GACATCCTGATGACCCAATCTCCATCCTCCATGTCTGTATCTCTGGGAGACACAGT<br>CAGCATCACTTGCCATGCAAGTCAGGGCATTGGCAGTAATATAGGGTGGTTGCAG<br>CAGAAACCAGGGAAATCATTTAAGGGCCTGATCTATCTTGGAACCAACTTGGAAGA<br>TGGAGTTCCATCAAGGTTCAGTGGCAGTGGATCTGGAGCAGATTATTCTCTCACC<br>ATCAGCAGCCTGGAATCTGAAGATTTTGCAGACTATTACTGTGTACAGTATGTTCA<br>GTTTCCGTACACGTTCGGAGGGGGGACCAAGCTGGAAATAAAA |
| Ab17 | GAGGTCCAGCTGCAGCAGTCTGGACCTGAGCTGGTGAAGCCTGGGGCTTCAGT<br>GAAGATATCCTGCAAGGCTTCTGGTTACTCATTCACTGACTACTACATGAACTGGG<br>TGAAGCAAAGTCCTGAAAAGAGCCTTGAGTGGATTGGAGAGATTAATCCTAACACT<br>GGTGGTACTACCTACAACCAGAAGTTCAAGGCCAAGGCCACATTGACTGTAGACA<br>AATCCTCCAGCACAGCCTACATGCAGCTCAAGAGCCTGACATCTGAGGACTCTGC<br>AGTCTATTACTGTGCAAGATACTATGGTAACCTCTATGCTATGGACTACTGGGGTC<br>AAGGAACCTCAGTCACCGTCTCCTCA | GACATCCTGATGACCCAATCTCCATCCTCCATGTCTGTATCTCTGGGAGACACAGT<br>CAGCATCACATGCCATGCAAGTCAGGGCATAAGTAGTAATATAGGGTGGTTGCAG<br>CAGAAACCAGGGAAATCATTTAAGGGCCTGATCTATCATGGAACCAACTTGGAAGA<br>TGGAGTTCCATCAAGGTTCAGTGGCAGTGGATCTGGAGCAGATTATTCTCTCACC<br>ATCAGCAGCCTGGAATCTGAAGATTTTGCAGACTATTACTGTGTACAGTATGTTCA<br>GTTTCCGTACACGCTCGGAGGGGGGACCAAGCTGGAAATAAAA |
| Ab19 | CAGGTTCAGCTGCAGCAGTCTGGAGCTGAGCTGGCGAGGCCTGGGGCTTCAGT<br>GAAGCTGTCCTGCAAGGCTTCTGGCTACCCCTTCACAAGCTATGGTATAAACTGG<br>GTGAAGCAGAGAACTGGACAGGGCCTTGAGTGGATTGGAGAGATTTATCCTAGAA<br>TTGGAAATACTTACTATAATGAGAAGTTCAAGGGCAAGGCCACACTGACTGCAGAC<br>AAATCCTCCAGCACAGCGTACATGGAGTTCCGCAGCCTGACATCTGAGGACTCTG<br>CGGTCTATTTCTGTGCAAGATCGTGGAATAGTAACTACGGGGAGTACTACTTTGA<br>CTACTGGGGCCAAGGCACCACTCTCACAGTCTCC | GACATTGTGATGACCCAGTCTCACAAATTCATGTCCACATCAATAGGAGACAGGGT<br>CAGCATCACCTGCAAGGCCAGTCACGATGTGAGTACTGCTGTAGCCTGGTATCAA<br>CAAAAACCAGGGCAATCTCCTAAGTTACTGATTTACTGGGCATCCACCCGGCACAC<br>TGGAGTCCCTGATCGCTTCACAGGCAGTGGATCTGGGACAGATTATACTCTCACC<br>ATTAGAAGTGTGCAGGCAGAAGACCTGGCACTTTATTACTGTCAGCAACATTATAG<br>CACTCCGTACACGTTCGGAGGGGGGACCAAGCTGGAAATAAAA |
| Ab20 | GAGGTCCAGCTGCAGCAGTCTGGACCTGAGCTGGTGAAGCCTGGGGCTTCAGT<br>GAAGATATCCTGCAAGGCTTCTGGTTTCTCATTCACTGGCTACTCCATGAACTGGA<br>TGAAACAAAGTCCTGAAAAGAGCCTTGAGTGGATTGGAGAAATTAATCCTACCACT<br>GGTGGTACTACCTACAACCAGAAGTTCAAGGCCAAGGCCACATTGACTGTAGACA<br>AATCCTCCAGCACAGCCTACATACAACTCAAGAGCCTGACATCTGAGGACTCTGCA<br>GTCTATTACTGTGCAAGGGGCCGGGCCGACTACTGGGGCCAAGGCACCACTCTC<br>ACAGTCTCCTCA | GACATTGTGCTCACCCAATCTCCAGCTTCTTTGGCTGTGTCTCTAGGGCAGAGAG<br>CCACCATCTCCTGCAGAGCCAGTGAAAGTGTTGAATATTATGGCACAGGTTTAGT<br>GCAGTGGTTCCAACAGAAACCAGGACAGCCACCCAAACTCCTCATCTATGCTGCC<br>TCCAACGTGGAATCTGGGGTCCCTGCCAGGTTTAGTGGCAGTGGGTCTGGGACA<br>GACTTCAGCCTCAACATCCATTCTGTGGAGGAGGATGATATTGCAATGTATTTCTG<br>TCACCAAAGTAGGAAGCTTCCGTGGACGTTCGGTGGAGGCACCAAGCTGGAAAT<br>CAAA |

**Table S1.** VH and VL sequences for antibodies selected for recombinant expression and characterization as shown in **Fig. 4E.**
